# Protein secretion by the type IV pilus machinery in *Francisella tularensis*

**DOI:** 10.64898/2026.05.11.724269

**Authors:** Angela DeRosa, P. Todd Benziger, Vinaya Sampath, Erik J. Kopping, David G. Thanassi

## Abstract

*Francisella tularensis* is a highly virulent, Gram-negative bacterial pathogen that causes the zoonotic disease tularemia. *F. tularensis* infects a variety of host cells and replicates intracellularly while evading and interfering with host immune responses. The molecular mechanisms that facilitate the intracellular replication and virulence of *F. tularensis* are poorly understood. The *Francisella* genome contains a set of *pil* genes that code for the assembly of surface fibers termed type IV pili (T4P). T4P are major bacterial virulence determinants but the function of the *pil* system during *F. tularensis* infection and intracellular growth is unclear. T4P are closely related to the type II secretion pathway and the *pil* system of a related *Francisella* species, *F. novicida*, was shown to function in protein secretion as well as pilus assembly. To identify proteins secreted by *F. tularensis*, we analyzed the *F. tularensis* Live Vaccine Strain (LVS) using bio-orthogonal non-canonical amino acid tagging (BONCAT). Using BONCAT in conjunction with proteomics, we identified candidate proteins secreted by the wild-type LVS, as well as candidate proteins whose extracellular abundance decreased in the absence of the PilF ATPase or the PilE4 pilus subunit. Using epitope tagging of selected candidates, we validated T4P-mediated secretion of the ChiA and ChiD chitinases and the KatG catalase by the LVS. These results further our understanding of the *pil* system and protein secretion pathways in *F. tularensis*.

**IMPORTANCE:** *Francisella tularensis* is a highly virulent Gram-negative bacterial pathogen and the causative agent of tularemia. *F. tularensis* lacks secretion systems utilized by other intracellular bacterial pathogens but contains *pil* genes that encode for type IV pili (T4P) and may also function in protein secretion. T4P are observed on the surface of all *Francisella* spp. but *pil*-mediated protein secretion has only been reported for *F. novicida*, which is not normally pathogenic in humans. In this study, we used bio-orthogonal non-canonical amino acid tagging to identify proteins secreted by *F. tularensis*, for which there is limited information. We demonstrate that the *F. tularensis <u>pil</u>* system is capable of protein secretion and validate T4P-medeated secretion of the ChiA and ChiD chitinases and the KatG catalase. These results will facilitate investigation of *Francisella* virulence mechanisms and may provide targets for therapeutic intervention.

## INTRODUCTION

*Francisella tularensis* is a Gram-negative bacterium that causes tularemia, a severe and potentially fatal infection in humans (1, 2). *F. tularensis* is one of the most infectious bacteria known, with 10 or fewer organisms sufficient to cause a life threatening illness. *F. tularensis* is a zoonotic pathogen with the ability to adapt and thrive in diverse environments and within a broad range of hosts, including arthropods and mammals. Transmission to humans occurs via various routes, including direct contact with bacteria, bites from arthropod vectors, ingestion of contaminated materials, or inhalation of aerosolized organisms (1, 2). Due to its low infectious dose, ease of aerosolization, and historical use as a bioweapon, *F. tularensis* is classified as a Tier 1 Select Agent by the Centers for Disease Control and Prevention.

There are two clinically relevant species of *F. tularensis* (1, 2). *F. tularensis* subsp. *tularensis* is the “type A” biovar that causes the most severe form of tularemia, leading to high mortality rates without immediate therapeutic intervention. The “type B” biovar, *F. tularensis* subsp. *holarctica*, produces a milder disease in humans. The *F. tularensis* Live Vaccine Strain (LVS) was generated by serial passage of a subsp. *holarctica* strain. The LVS is attenuated for virulence in humans but remains pathogenic in animals such as mice, where it closely mimics infection with fully virulent *F. tularensis* and provides a model for the human disease that can be studied under biosafety level 2 (BSL2) conditions. An additional *Francisella* species, *F. novicida*, is closely related to *F. tularensis* and has proven useful as a model strain that can also be studied under BSL2 conditions (3). However, *F. novicida* is considered a more environmentally adapted strain and exhibits differences from *F. tularensis* in cell culture and mouse infection models (4).

*F. tularensis* is a facultative intracellular pathogen that can infect and replicate within a wide variety of host cells, with the macrophage being a preferred replicative niche *in vivo* (5). During infection, the bacteria evade and suppress the host’s immune response to promote their survival and replication within the host cell cytosol (6–8). Following adherence to and uptake by a macrophage, *F. tularensis* is initially confined within the *Francisella-*containing phagosome. The bacteria delay phagosomal maturation, resist host antimicrobial defenses, and rapidly escape to the host cell cytosol (9, 10). Escape from the *Francisella-*containing phagosome happens within 1-4 hours of initial uptake and is dependent on genes encoded within the *Francisella* pathogenicity island (FPI) (11, 12). The evasion and manipulation of host defense mechanisms, together with a metabolism optimized for cytosolic replication, allows *F. tularensis* to preserve its intracellular niche and replicate to high numbers prior to bacterial escape from the host cell and systemic spread (13, 14). Survival and replication within host cells typically requires the delivery of effector proteins via bacterial secretion systems. *Francisella* spp. lack the type III and type IV secretion systems that are often used by intracellular Gram-negative bacteria to deliver effector proteins into host cells (15, 16). However, the FPI codes for a type VI secretion system that is critical for bacterial escape from the phagosome (17). Other secretion systems present in *Francisella* include a type I secretion system and the production of outer membrane vesicles and tubes (OMVT) (18–20). In addition, the *Francisella* genome contains a set of *pil* genes that code for the assembly of type IV pili (T4P) and are related to bacterial type II secretion systems (T2SS) (21–23).

T4P are multifunctional, hairlike surface fibers expressed by a wide range of bacteria, with roles in motility, adhesion to surfaces, biofilm formation, and host-pathogen interactions (24, 25). T4P have been observed on the surface of *F. tularensis subsp. tularensis, F. tularensis subsp. holartica* LVS, and *F. novicida* (23, 26–28). The T4P contribute to *Francisella* host cell adhesion and virulence, through mechanisms that are yet to be characterized (26, 27, 29, 30). In *F. novicida*, the *pil* system was also shown to contribute to the secretion of extracellular proteins in addition to the assembly of T4P (28, 30). The *F. novicida* secreted proteins include two chitinases (ChiA and ChiB), a chitin binding protein (CbpA), a protease (PepO), a ß-glucosidase (BglX), and two proteins of unknown function (Fsp53 and Fsp58) (30).

Core components of the T4P machinery include an inner membrane pilus assembly platform (PilC) and an outer membrane channel termed a secretin (PilQ) (Fig 1A). In typical T4P systems, polymerization of pilus subunit proteins (pilins) into the pilus fiber is driven by an assembly ATPase and disassembly of the fiber is driven by a retraction ATPase. In *Francisella*, PilF and PilT are thought to represent the assembly and disassembly ATPases, respectively.

**Figure 1.**
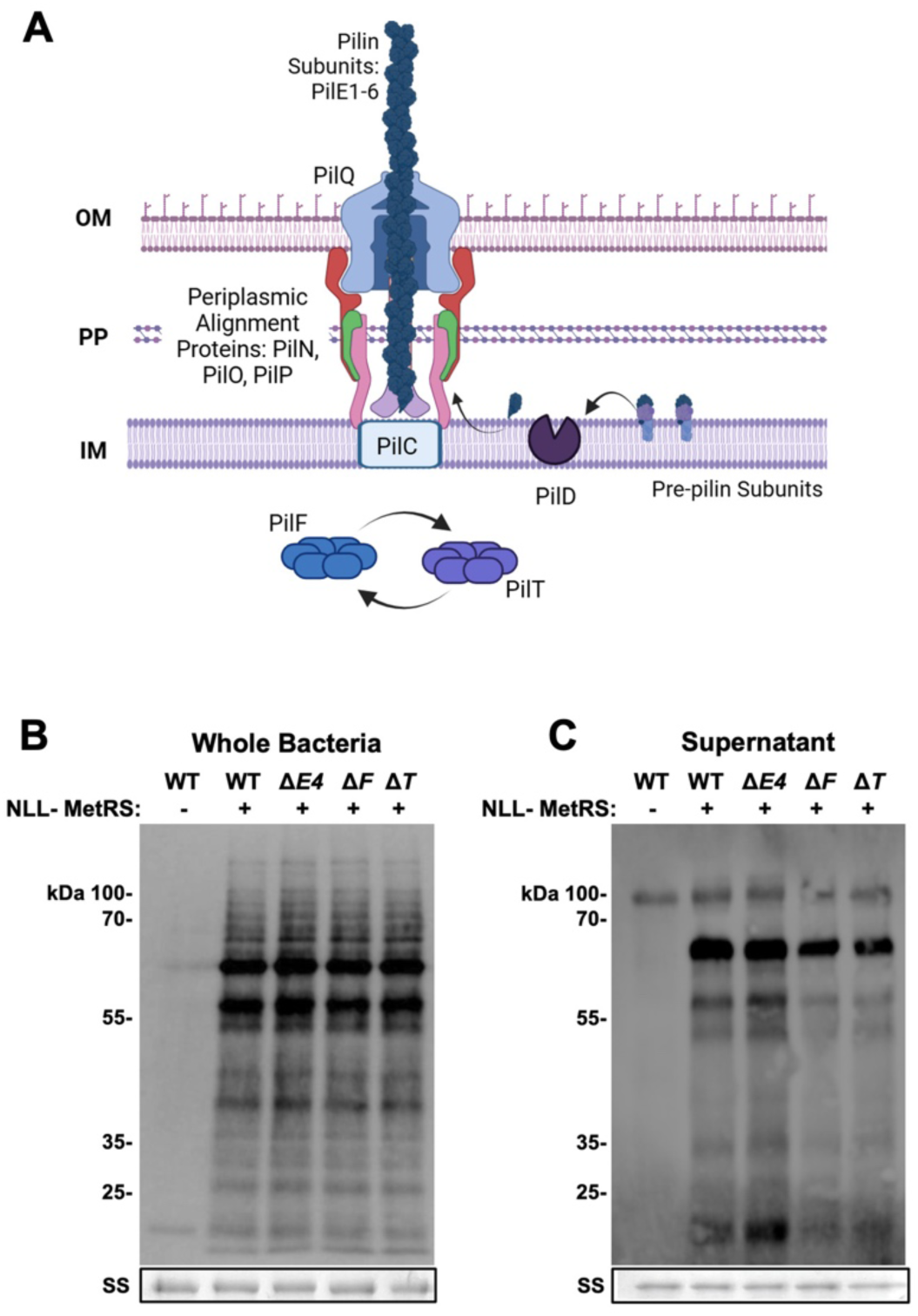
BONCAT labeling of proteins secreted by the LVS. **(A)** Overview of the T4P system in *F. tularensis*. Pilus subunits (PilE) are matured by the PilD pre-pilin peptidase and assembled into the pilus fiber at the PilC assembly platform. PilE4 is thought to be the major pilin subunit. PilF and PilT are cytoplasmic ATPases that, in canonical systems, energize pilus assembly and retraction, respectively. Protein and pilus fiber secretion occurs through the PilQ OM channel. **(B and C)** The LVS WT and Δ*pilE4* (Δ*E4*), Δ*pilF* (Δ*F*), and Δ*pilT* (Δ*T*) mutant strains, without (-) or with (+) the *att*::NLL-MetRS tRNA synthase were incubated in minimal infection medium in the presence of ANL. Whole bacterial lysates **(B)** or cell-free supernatant fractions **(C)** were collected, labeled with biotin via click chemistry, and analyzed by SDS-PAGE and blotting using streptavidin-HRP. Parallel samples were detected by silver staining (SS) and sections from these gels are shown as loading controls (boxed rows). The images shown are representative of three independent experiments.

PilF is required for T4P formation and is also required for *pil*-dependent protein secretion in *F. novicida* (27, 28, 30). Distinct from canonical T4P systems, it is unclear whether the *Francisella* T4P retract, and pilus fibers are lost when PilT is mutated, in contrast to the hyper-piliation phenotype that is seen in other T4P systems (27, 28). Loss of PilT diminishes protein secretion in *F. novicida* but secretion is not dependent on PilT (28, 30). The *pilT* genes in *F. novicida* and *F. tularensis* subsp. *tularensis* strains are intact (31, 32). However, in the LVS and other *F. tularensis* subsp. *holarctica* strains, *pilT* contains a point mutation that creates a premature stop codon and splits the gene into two open reading frames. There is contradictory evidence for whether there is read-through of the stop codon to produce a full-length PilT protein in the LVS (21, 23, 27).

The T4P fiber is typically composed of a major pilin and a set of additional minor pilins that are involved in pilus assembly and function (Fig 1A). The *Francisella* genome encodes six potential pilin subunits, PilE1-6, although there is variation in both the number of pilin genes present and their coding sequences among strains (22, 23, 26, 33, 34). PilE4 is thought to be the major pilus structural subunit, as deletion of *pilE4* eliminates pilus expression, whereas mutations in the other pilin genes do not (26, 28). Of note, the *pilE1-3* genes in the LVS are non-functional due to deletions and truncations of their genomic regions, with impacts on virulence (23, 29). In *F. novicida*, distinct roles have been assigned to PilE1 and PilE4, with PilE1 involved in protein secretion and PilE4 important for T4P fiber formation (28, 30). This raises the question of whether the *F. tularensis* LVS is capable of protein secretion, given that its *pilE1* gene is mutated.

In this study, we used bio-orthogonal non-canonical amino acid tagging (BONCAT) to examine protein secretion in *F. tularensis*, identifying proteins secreted by the bacteria under conditions used for the infection of mammalian cells. Comparison of proteins secreted by the wild-type (WT) LVS with Δ*pilE4*, Δ*pilF*, and Δ*pilT* mutants identified secreted proteins whose extracellular abundance was significantly decreased in both PilF and PilT mutants. We found PilE4 to have a more limited effect on protein secretion, consistent with its proposed role as the major T4P subunit. Complementation of the Δ*pilF* and Δ*pilE4* mutants restored protein secretion into the supernatant; however, complementation of the Δ*pilT* mutant, using either the LVS or *F. novicida* sequence, was unable to rescue protein secretion. Therefore, the function of PilT in the LVS remains unclear. Overall, these results demonstrate the presence of T4P-mediated protein secretion in *F. tularensis* and confirm that the system functions as a hybrid pilus assembly and protein secretion machinery. Improved understanding of the T4P system and the identification of secreted proteins will provide insights into *F. tularensis* virulence mechanisms and the pathogenesis of tularemia.

## RESULTS

### Identification of proteins secreted by *F. tularensis* using BONCAT

The BONCAT method allows for the selective labeling and detection of bacterial proteins, providing a powerful approach to identify secreted proteins in complex environments, such as during infection of host cells (35). We also found this method to be optimal for detecting proteins secreted by *F. tularensis* in simpler environments, such as *in vitro* culture in the absence of host cells. In the BONCAT method, bacteria express a modified methionyl-tRNA synthetase that enables the incorporation of a non-canonical amino acid, azidonorleucine (ANL), into newly synthesized bacterial proteins at methionine codons, allowing for subsequent labeling of the proteins with a biotin-alkyne tag via click chemistry (35). The biotinylated proteins are then detected or purified using streptavidin and protein identities determined by mass spectrometry. To use BONCAT in *F. tularensis*, a codon-optimized NLL-MetRS tRNA synthase was inserted into the chromosomal *att* site of the LVS, generating strain LVS *att*::NLL-MetRS.

To confirm the selective labeling and detection of *F. tularensis* proteins by BONCAT, the LVS *att*::NLL-MetRS and parental WT strain were grown to mid-log phase and then shifted for 4 h to a minimal cell culture medium, in the presence of ANL, to mimic an infection setting. Analysis of whole bacterial lysates by blotting with streptavidin-HRP showed a broad labeling of proteins in the *att*::NLL-MetRS strain and an almost complete absence of labeling in the WT LVS (Fig. 1B). Analysis of cell-free supernatant fractions showed a similar selective labeling of proteins in the *att*::NLL-MetRS strain (Fig. 1C), confirming the ability of BONCAT to detect *F. tularensis* secreted proteins. A prominent non-specific band at ∼100 kDa was present in the supernatant fraction of the WT LVS, likely representing a naturally biotinylated protein (Fig. 1C). The pattern of labeled secreted proteins in the LVS *att*::NLL-MetRS was distinct from that detected in whole bacteria, indicating that the proteins were not simply released by bacterial lysis. Prominent labeled bands were present above the 55 kDa marker in both whole bacteria and the supernatant fraction, although whether these are the same or distinct proteins remains to be determined (Fig. 1B and C). A prominent low molecular mass band (∼18 kDa) was only detected in the cell-free supernatant fraction (Fig. 1).

To identify proteins secreted by the LVS, cell-free supernatant fractions of the LVS *att*::NLL-MetRS grown in the presence of ANL as described above were collected, labeled with biotin, and then purified using streptavidin magnetic beads for analysis by mass spectrometry. Mass spectrometry identified >400 proteins present in the supernatant fraction (Table S1). Table 1 shows the 25 most abundant proteins detected. The proteins include many that were previously identified as secreted by *Francisella*, supporting our approach (19, 20, 36–39). The KatG catalase was the sixth most abundant protein detected. KatG is an important *F. tularensis* secreted virulence factor, allowing the bacteria to counteract oxidative stress encountered during host cell infection (40). The KatG secretion mechanism is incompletely understood, although it is a prominent constituent of OMVT released by the bacteria (19, 20, 41, 42). Several additional of the identified proteins are also present in OMVT, including OmpH and SucC (Table S1) (19, 41). Additional members of the identified proteins are associated with the *Francisella* T6SS, including IglB and ClpB (Table 1) (17, 36). Other proteins identified by our BONCAT analysis are thought to be primarily cytoplasmic, including abundant proteins that are often detected in mass spectrometry analyses.

**Table 1.** Top 25 must abundant LVS secreted proteins identified by BONCAT.

| Average Abundance | Locus Tag | Gene Symbol | Description | MW [kDa] | Pfam IDs |
| --- | --- | --- | --- | --- | --- |
| 2.58E+09 | FTL_0094 | clpB | Heat-shock protein, chaperone | 96 | Pf00004, Pf02861, Pf07724, Pf10431, Pf17871 |
| 1.09E+09 | FTL_0519 | minD | Septum site-determining protein | 30.1 | Pf13614 |
| 8.73E+08 | FTL_1041 |  | diphosphate synthase | 36.4 | Pf00348 |
| 4.58E+08 | FTL_1714 | groEL | Heat-shock protein, chaperone | 57.4 | Pf00118 |
| 4.52E+08 | FTL_1751 | tuf | Elongation factor (EF-Tu) | 43.4 | Pf00009, Pf03143, Pf03144 |
| 3.90E+08 | FTL_1504 | katG | Catalase-peroxidase | 82.5 | Pf00141 |
| 2.89E+08 | FTL_1191 | dnaK | Heat-shock protein, chaperone | 69.1 | Pf00012 |
| 2.67E+08 | FTL_0653 |  | Sulfur acceptor protein, SufE | 15.7 | Pf02657 |
| 2.32E+08 | FTL_0009 |  | Outer membrane protein of unknown function | 19.5 | Pf03938 |
| 1.59E+08 | FTL_0267 | htpG | Heat shock protein, chaperone, Hsp90 | 72.3 | Pf00183, Pf13589 |
| 1.03E+08 | FTL_0234 | fusA | Elongation factor (EF-G) | 77.7 | Pf00009, Pf00679, Pf03144, Pf03764, Pf14492 |
| 9.29E+07 | FTL_1158 | iglB | Intracellular growth locus protein B | 57.9 | Pf05943, Pf18945 |
| 8.04E+07 | FTL_1060 |  | Carboxypeptidase | 48 | Pf00768, Pf07943 |
| 7.55E+07 | FTL_1478 | guaB | Dehydrogenase/GMP reductase | 52.1 | Pf00478, Pf00571 |
| 7.13E+07 | FTL_0546 |  | Diphosphate synthase/farnesyl diphosphate synthase | 32.8 | Pf00348 |
| 7.04E+07 | FTL_0572 |  | Protein of unknown function | 51.9 | Pf20891 |
| 6.96E+07 | FTL_0573 |  | Protein of unknown function | 35.8 | Pf13036 |
| 6.84E+07 | FTL_1916 |  | Competence protein | 76.4 | Pf03772, Pf13567 |
| 6.77E+07 | FTL_1908 | ftsA | Cell division protein | 44.8 | Pf02491, Pf14450 |
| 6.50E+07 | FTL_0269 | gdh | Glutamate dehydrogenase | 49.1 | Pf00208, Pf02812 |
| 6.28E+07 | FTL_1743 |  | RNA polymerase, beta' subunit | 157.3 |  |
| 6.15E+07 | FTL_0298 |  | Protein of unknown function | 74.2 |  |
| 6.09E+07 | FTL_1592 | accB | Acetyl-CoA carboxylase, biotin carboxy carrier protein subunit | 16.4 | Pf00364 |
| 6.04E+07 | FTL_1907 | ftsZ | Cell division protein | 39.7 | Pf00091, Pf12327 |
| 6.02E+07 | FTL_1225 |  | Protein of unknown function | 25.7 |  |

### Protein secretion is altered in *F. tularensis pil* mutants

To investigate involvement of the T4P system in protein secretion by *F. tularensis*, we next used BONCAT to compare the secretion profiles of the WT LVS with Δ*pilE4*, Δ*pilF*, and Δ*pilT* deletion mutant strains. PilE4 is the putative major pilin subunit, but it is not thought to be involved in protein secretion (26, 30). PilF is the T4P assembly ATPase and was previously shown to be required for both pilus assembly and protein secretion (27, 28, 30). PilT is the putative T4P retraction ATPase but its role in *Francisella* is unclear (27, 30). For BONCAT analysis, the NLL-MetRS tRNA synthetase was inserted into the chromosome of each of the mutants as done for the WT LVS. No growth differences were observed among the engineered WT and mutant LVS strains (Fig. S1).

Analysis of whole bacterial lysates by blotting with streptavidin-HRP revealed a similar profile of labeled proteins for the WT and mutant strains (Fig. 1B), indicating that the *pil* deletion mutants did not affect general protein synthesis or ANL incorporation. Analysis of the cell-free supernatant fractions for the presence of secreted proteins showed that the LVS Δ*pilE4 att*::NLL-MetRS mutant had a similar profile to the parental WT strain (Fig. 1C), indicating that PilE4 does not play a major role in protein secretion, as expected. In contrast, reductions in the intensity of labeled bands were observed for the LVS Δ*pilF* and Δ*pilT att*::NLL-MetRS mutants (Fig. 1C). Based on previous studies, it was expected that PilF would mediate protein secretion but a role for PilT was unexpected (28, 30). Overall, these results suggest an important role for the *F. tularensis* T4P system in protein secretion.

### ChiA, ChiD, and KatG are secreted by the T4P system in *F. tularensis*

To identify potential T4P-dependent secreted proteins, BONCAT labeled proteins in the cell-free supernatant fractions of the LVS WT, Δ*pilF*, and Δ*pilE4 att*::NLL-MetRS strains were collected and analyzed by mass spectrometry as described above. Comparison of the WT and mutant strains indicated many proteins with altered abundance in the Δ*pilF* and Δ*pilE4* mutants, consistent with the results shown in Fig. 1C and supporting a role for the T4P system in protein secretion in *F. tularensis*. However, variability in the BONCAT data prevented conclusive identification of *pilF*-or *pilE*-dependent secreted proteins. We therefore selected several candidate T4P-dependent secreted proteins for follow-up analysis based on the BONCAT results, previous evidence for secretion, and potential roles as effector proteins. We chose the following proteins for analysis: the ChiA (FTL_1521) and ChiD (FTL_1793) chitinases; the catalase KatG (FTL_1504); the SucC synthetase (FTL_1553); the acetyl-CoA carboxylase, biotin carboxyl carrier protein AccB (FTL_1592); and the hypothetical proteins FTL_0569 and FTL_1225.

Genes encoding each of the candidate proteins with a triple FLAG-tag appended to their C termini were inserted into the chromosomal *att* site of the *F. tularensis* LVS WT and Δ*pilE4*, Δ*pilF*, and Δ*pilT* deletion mutant strains. Bacteria expressing the tagged proteins under their native promoters were then incubated in infection medium as above and whole bacterial lysates and cell-free supernatant fractions were collected and analyzed for the presence of the tagged proteins by blotting with anti-FLAG antibodies. Each of the tagged proteins was detected in the whole bacterial lysates (Figs. 2 and 3). The protein levels were comparable between the WT and deletion mutant strains, indicating that loss of PilE4, PilF, and PilT did not affect expression of the tagged proteins. Each of the tagged proteins was also detected in the cell-free supernatant fractions of the WT LVS bacteria (Figs. 2 and 3). This confirms that the proteins are indeed secreted by *F. tularensis* into the extracellular environment.

**Figure 2.**
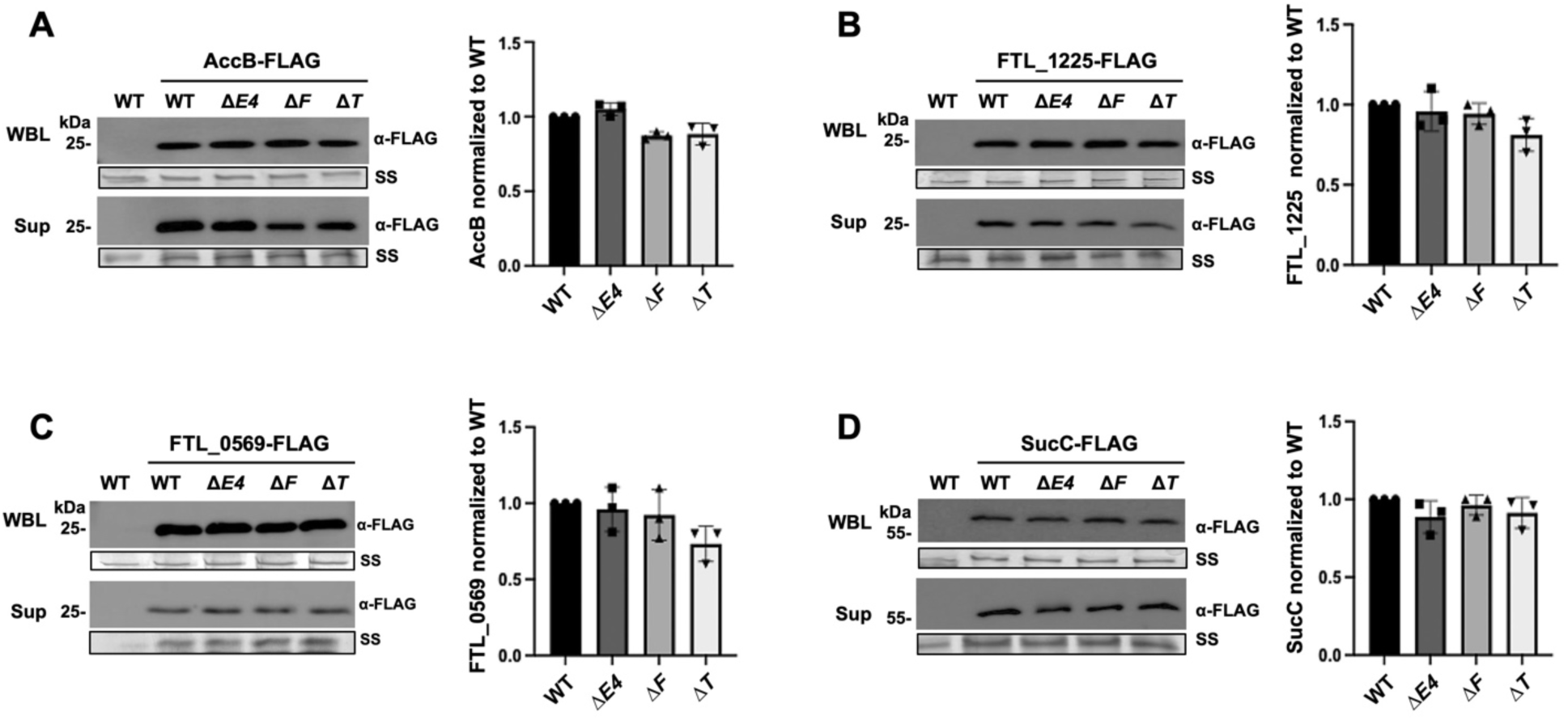
Analysis of FLAG-tagged candidates reveals proteins secreted independently of the T4P system. Whole bacterial lysates (WBL) and cell-free supernatant fractions (Sup) were collected from the LVS WT and Δ*pilE4* (Δ*E4*), Δ*pilF* (Δ*F*), and Δ*pilT* (Δ*T*) mutant strains following incubation in minimal infection medium. The strains expressed FLAG-tagged versions of AccB **(A)**, FTL_1225 **(B)**, FTL_0569 **(C)**, or SucC **(D)**. The parental WT strain was used as a negative control. The samples were analyzed by SDS-PAGE and blotting with anti-FLAG antibodies. Parallel samples were detected by silver staining (SS) and sections from these gels are shown as loading controls (boxed rows). The graph to the right in each panel shows densitometry analysis of the bands in the corresponding supernatant fraction, normalized to the WT LVS. Data represent means ± standard deviation (SD) from three independent experiments. No significant differences were observed; calculated by one-way ANOVA with Brown-Forsythe test and pairwise t-test.

**Figure 3.**
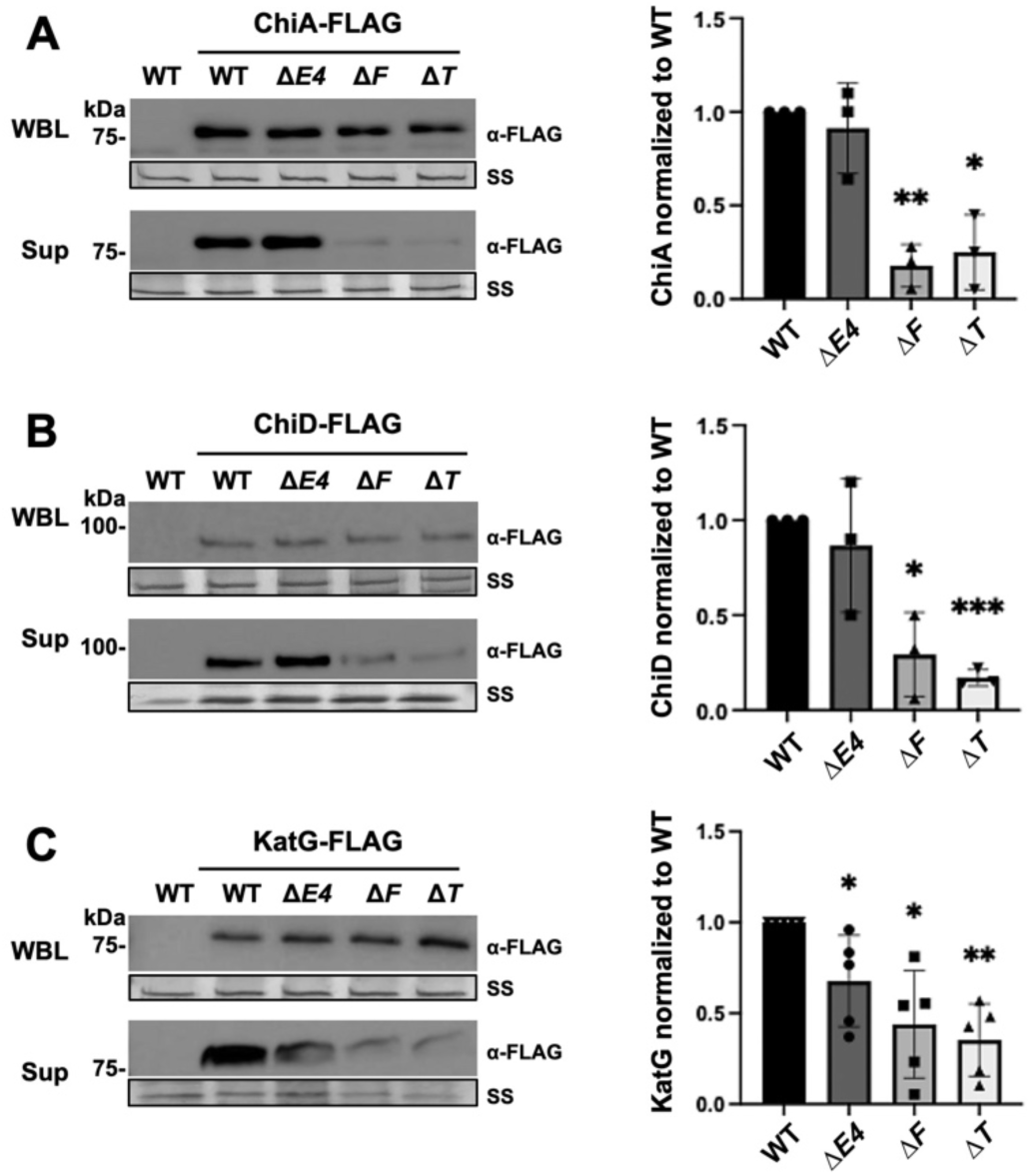
Analysis of FLAG-tagged candidates reveals proteins dependent on the T4P system for secretion. Whole bacterial lysates (WBL) and cell-free supernatant fractions (Sup) were collected from the LVS WT and Δ*pilE4* (Δ*E4*), Δ*pilF* (Δ*F*), and Δ*pilT* (Δ*T*) mutant strains following incubation in minimal infection medium. The strains expressed FLAG-tagged versions of ChiA **(A)**, ChiD **(B)**, or KatG **(C)**. The parental WT strain was used as a negative control. The samples were analyzed by SDS-PAGE and blotting with anti-FLAG antibodies. Parallel samples were detected by silver staining (SS) and sections from these gels are shown as loading controls (boxed rows). The graph to the right in each panel shows densitometry analysis of the bands in the corresponding supernatant fraction, normalized to the WT LVS. Data represent means ± SD from three independent experiments. *, *P* < 0.05; **, *P* < 0.01; ***, *P* < 0.001; calculated by one-way ANOVA with Brown-Forsythe test and pairwise t-test.

Comparison of the cell-free supernatant fractions of the LVS WT, Δ*pilE4*, Δ*pilF*, and Δ*pilT* strains revealed that the AccB, FTL_1225, FTL_0569, and SucC proteins were detected at equivalent levels in supernatant fractions from the WT and deletion mutant bacteria (Fig. 2). Thus, these four proteins are secreted by *F. tularensis* in a T4P independent manner. In contrast, extracellular levels of the ChiA, ChiD, and KatG proteins were each decreased in the *pil* mutant strains (Fig. 3). ChiA, ChiD, and KatG levels in the supernatant fraction were decreased in the Δ*pilF* and Δ*pilT* strains. The secretion of KatG, but not ChiA or ChiD, was also decreased in the supernatant fraction of the Δ*pilE4* mutant, although the KatG secretion defect in the Δ*pilE4* mutant was not as severe as for the Δ*pilF* and Δ*pilT* mutants (Fig. 3). These data confirm that *F. tularensis* uses the T4P system for protein secretion. Further, the results suggest a non-canonical role for PilT in protein secretion in the LVS and indicate that the putative major pilin subunit, PilE4, functions in secretion for at least one protein, KatG.

The levels of secreted ChiA, ChiD, and KatG were greatly reduced in the LVS Δ*pilF* and Δ*pilT* strains, but not completely absent. This remaining low level could be due to secretion by another pathway or background lysis of the bacteria. Both ChiA and KatG have been detected in OMVT released by *Francisella* spp. (19, 20, 41). To test if OMVT account for the remaining secretion observed in the mutants, cell-free supernatant fractions from the LVS WT, Δ*pilF*, and Δ*pilT* strains expressing FLAG-tagged ChiA were subjected to ultracentrifugation to remove any vesicles present. For each of the strains, pelleting out the OMVT did not alter the levels of secreted ChiA detected (Fig. S2). We further blotted the supernatant fractions using an antibody to the FopA outer membrane protein, both as a marker for OMVT (41) and to detect bacterial lysis. No FopA was detected in the supernatant fractions from the WT or mutant strains (Fig. S2). These results suggest that neither OMVT nor background bacterial lysis is responsible for the low level of secreted proteins remaining in the LVS Δ*pilF* and Δ*pilT* mutants.

To confirm the secretion phenotypes of ChiA, ChiD, and KatG in the Δ*pilE4*, Δ*pilF*, and Δ*pilT* mutant backgrounds, we complemented the mutants with plasmids expressing the deleted gene under its native promoter. As shown in Fig. 4, complementation of the Δ*pilF* mutant increased ChiA, ChiD, and KatG secretion into the cell-free bacterial supernatant fraction.

**Figure 4.**
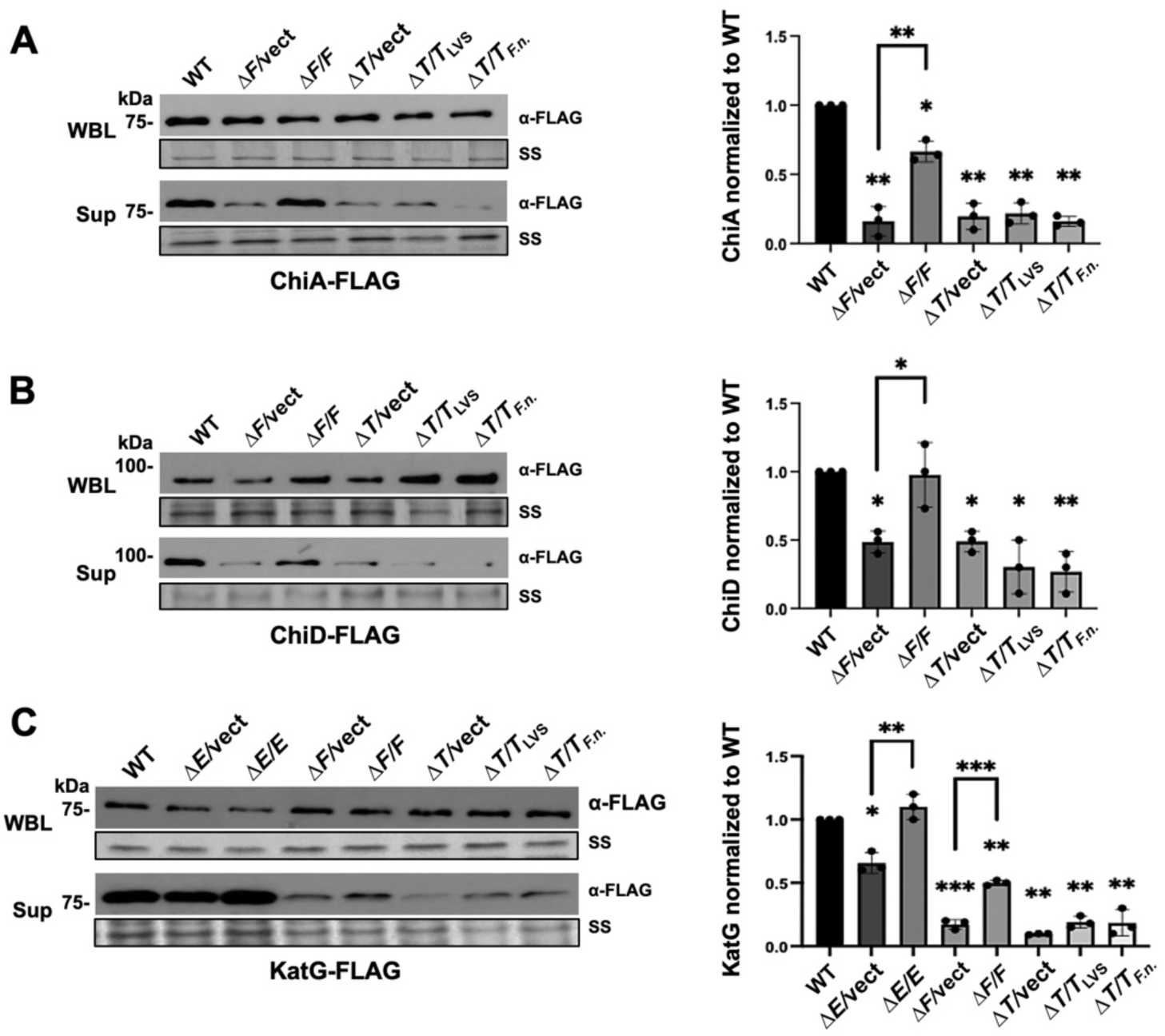
Complementation analysis of the LVS *pil* deletion mutants. Whole bacterial lysates (WBL) and cell-free supernatant fractions (Sup) were collected from the LVS WT and Δ*pilE4* (Δ*E4*), Δ*pilF* (Δ*F*), and Δ*pilT* (Δ*T*) mutant strains following incubation in minimal infection medium. The strains expressed FLAG-tagged versions of ChiA **(A)**, ChiD **(B)**, or KatG **(C)**. The mutant strains were complemented with plasmids expressing the deleted gene, or transformed with vector only (vect) as a negative control. The *pilT* gene from the LVS or *F. novicida* was used for complementation of the Δ*pilT* mutant strain. The samples were analyzed by SDS-PAGE and blotting with anti-FLAG antibodies. Parallel samples were detected by silver staining (SS) and sections from these gels are shown as loading controls (boxed rows). The graph to the right in each panel shows densitometry analysis of the bands in the corresponding supernatant fraction, normalized to the WT LVS. Data represent means ± SD from three independent experiments. *, *P* < 0.05; **, *P* < 0.01; ***, *P* < 0.001; calculated by one-way ANOVA with Brown-Forsythe test and pairwise t-test.

Complementation of the Δ*pilE4* mutant also rescued secretion of KatG (Fig. 4C). However, we were unable to complement the secretion defects observed for the Δ*pilT* mutant (Fig. 4). The *pilT* gene in the LVS contains a point mutation that creates a premature stop codon and divides the gene into two open reading frames (23, 27). Although there is evidence for read-through of the stop codon (23, 27), we reasoned that expression of the LVS *pilT* from a plasmid might not allow for complementation. The *F. novicida pilT* gene lacks the premature stop codon mutation present in the LVS but otherwise shares a high level of sequence homology to the LVS *pilT* (31, 32). Therefore, we tested complementing the LVS Δ*pilT* mutant with a plasmid expressing the *pilT* gene from *F. novicida* strain U112. However, the *F. novicida* PilT was also unable to complement the ChiA, ChiD, and KatG secretion defects observed in the LVS Δ*pilT* mutant (Fig. 4). Finally, we tested complementing the Δ*pilT* mutation with plasmids expressing the LVS or *F. novicida pilT* from a strong constitutive promoter. Although the over-expression plasmid caused increased levels of background proteins in the cell-free supernatant fraction, presumably due to bacterial lysis, we did not observe rescue of ChiA or ChiD secretion using these plasmids (Fig. S3). Based on our inability to complement the LVS Δ*pilT* mutation, the role of PilT in protein secretion by *F. tularensis* remains unclear.

## DISCUSSION

*F. tularensis* is a highly virulent intracellular pathogen with the ability to evade and subvert normal host defenses. Secreted effector proteins are expected to be key players in the ability of *F. tularensis* to infect host cells, replicate intracellularly, and cause disease. However, information is lacking regarding the identification of such effector proteins and their secretion pathways, particularly since *Francisella* spp. do not possess the type III and type IV secretion systems that are commonly used by Gram-negative intracellular pathogens to deliver virulence factors to host cells during infection (15, 16). In this study, we used BONCAT to examine proteins secreted by the *F. tularensis* LVS and identified three proteins that are secreted by the LVS T4P biogenesis system.

T4P are major virulence determinants for many bacterial species, with a range of functions including adhesion to host cells, colonization of surfaces, and motility (24, 25). T4P are closely related to the type II secretion pathway, and dual functions for the T4P machinery have been established in other Gram-negative pathogens. For example, the toxin co-regulated T4P system of *Vibrio cholerae* mediates secretion of a soluble colonization factor (43), and T4P in the ruminant foot rot pathogen *Dichelobacter nodosus* mediate secretion of extracellular proteases (44). T2SS are thought to function by loading a secreted cargo onto the tip of a pilus-like fiber in the periplasm, with extension of the fiber then mechanically pushing the substrate out through the channel of the OM secretin into the extracellular environment. Presumably, an analogous mechanism applies to T4P systems, where the assembling T4P fiber pushes a secreted substrate from the periplasm through the PilQ secretin to the extracellular environment.

T4P fibers have been observed on the surface of *F. tularensis* subsp. *holarctica* (the LVS), *F. tularensis* subsp. *tularensis* (strain Schu S4), and *F. novicida* (strain U112) (23, 26–28). There is evidence that the *Francisella* T4P function in host cell adhesion and virulence, but their specific contributions during infection remain unknown (26, 27, 29, 45). Previous studies in *F. novicida* demonstrated that the T4P system contributes to the secretion of soluble extracellular proteins in addition to the assembly of pilus fibers (28, 30). However, T4P-mediated protein secretion has not been characterized in *F. tularensis,* the causative agent most human infections. T4P fibers are composed of major and minor pilin subunits, which have multiple and diverse roles. The previous *F. novicida* analysis assigned distinct functions to the *Francisella* PilE1 and PilE4 pilins in protein secretion and pilus assembly, respectively (28, 30). PilE1 is absent in the LVS and therefore it was not clear that T4P-dependent protein secretion would be functional in the LVS, which could be part of the basis for its attenuation (23, 29). We report here that the LVS is indeed capable of T4P-mediated protein secretion, demonstrating that PilE1 is not required for this role in *F. tularensis*. Further, we found that PilE4, the putative major pilin, contributed to protein secretion in the LVS, although with a minor role compared to the PilF ATPase. These results highlight differences between *F. novicida* and *F. tularensis*. The *pilE1-E3* genes are intact in other *F. tularensis* strains, and it will be informative to determine how other *F. tularensis* strains compare with our findings in the LVS.

We used the BONCAT method to detect the global secreted protein profile of the *F. tularensis* LVS. BONCAT allows for the efficient labeling of newly synthesized bacterial proteins via click chemistry of a biotin alkyne tag and further detection or enrichment using streptavidin (35). This reduces the complexity of the sample and facilitates the identification of proteins of interest. In our exploration of different approaches, including mass spectrometry analysis of total cell-free supernatant fractions, we found BONCAT to be superior in allowing detection of secreted proteins and minimizing background interference (unpublished data). We also explored a variety of culture conditions to optimize protein secretion by the *F. tularensis* LVS during growth *in vitro*. We found that conditions used during bacterial infection of macrophages (even without the presence of host cells) allowed enhanced detection of proteins secreted into the extracellular medium. We hypothesize that the minimal cell culture medium utilized for these experiments causes an amino acid starvation response, which is also experienced during macrophage infection, thereby resulting in the increased production of secreted proteins (41, 46, 47).

Using BONCAT, we detected more than 400 candidate proteins secreted by the wild-type LVS under our culture conditions, including a variety of known proteins, previously identified virulence factors, and hypothetical proteins. Several of the candidate proteins were previously detected as constituents of membrane vesicles and tubes released by *Francisella*, including KatG, ChiA, and SucC (19, 20, 41). The T2SS has been shown in other bacteria to be involved in the delivery of secreted effector proteins to OMV for export to host cells (48). Our results suggest that the T4P system may perform a similar function in *F. tularensis*. The top secreted candidates also included proteins associated with the *F. tularensis* T6SS (ClpB and IglB), suggesting that the T6SS may be active under our culture conditions. Several of the secreted proteins identified in our analysis are more commonly associated with cytoplasmic functions.

The detection of highly abundant and stable cytoplasmic proteins is common in proteomics analyses and further studies are needed to demonstrate specific secretion versus non-specific release. However, some of these proteins were previously identified as secreted by *F. tularensis* and may have moonlighting functions during infection, as found in other bacteria (19, 20, 30, 36–39, 49).

Comparison of the BONCAT secretion profiles for the WT LVS versus the *pil* mutants indicated major contributions of the T4P system to extracellular protein secretion. Although the mass spectrometry analysis of the mutants was inconclusive due to variability in the data, we selected several candidate T4P-secreted proteins for further analysis: AccB, SucC, FTL_0569, FTL_1225, ChiA, ChiD, and KatG. Each of the candidate proteins was confirmed to be released by the LVS into the extracellular medium, but we found that AccB, SucC, FTL_0569 and FTL_1225 were released in a T4P-independent manner. AccB is a putative acetyl-CoA carboxylase biotin carboxyl carrier protein subunit (37). Enrichment of this protein in the BONCAT data may be due to its naturally biotinylated state. SucC (FTL_1533) is the succinyl-CoA ligase beta subunit involved substrate level phosphorylation in the TCA cycle. Its main function is carried out in the cytoplasm, but SucC has been reported in *Francisella* to be present in both membrane and surface locations, as well as present in OMVT (19, 37–39, 41). The hypothetical proteins FTL_0569 and FTL_1225 have appeared in LVS proteomic screens (37, 50, 51); FTL_0569 is a putative membrane protein with unknown function and FTL_1225 is thought to be an immunoreactive lipoprotein involved in the oxidative stress response. The mechanisms by which AccB, SucC, FTL_0569, FTL_1225 are released by *F. tularensis* into the extracellular medium remain to be defined.

Our epitope-tagging analysis confirmed that three of the candidate proteins, the chitinases ChiA and ChiD and the catalase KatG, are secreted from the LVS in a T4P-dependent manner. The presence of ChiA and ChiD in the extracellular medium was greatly diminished in the absence of the PilF ATPase. There are four chitinases encoded by *Francisella* spp.: ChiA, ChiB, ChiC, and ChiD, with differences in the encoding genes and chitinase activity among *Francisella* strains (52–54). The chitinases have been best characterized in *F. novicida*. Both ChiA and ChiB contribute to chitin colonization and biofilm formation, and the *F. novicida* chitinases also appear to modulate bacterial interactions with host cells (52, 55). ChiA and ChiB were previously found to be secreted by the T4P system in *F. novicida* (30). In comparison, we identified ChiA and ChiD as secreted by the *F. tularensis* LVS, but did not detect ChiB, revealing an additional distinction between *F. novicida* and *F. tularensis*. The chitinases have predicted Sec signal sequences and utilize the Sec system for delivery to the periplasm (52), where they could then engage the T4P system for secretion across the OM. Chitinases have established roles in environmental niches and vector colonization, but they are also emerging as effectors that contribute to colonization and infection of the mammalian host (56). Chitin binding proteins of *V. cholerae* enhance bacterial adherence to surface-exposed glycan moieties of human epithelial cells (57) and *Legionella pneumonia* chitinases promote bacterial survival in the lungs of infected mice (58). Additionally, *Salmonella enterica* serovar Typhimurium*, Serratia marcescens, Listeria monocytogenes,* and *Enterococcus faecalis* all utilize chitinases to promote bacterial attachment to and invasion of intestinal epithelial cells to encourage gastrointestinal infection (59–61). Further studies are needed to understand the contribution of the ChiA and ChiD chitinases to the virulence of *F. tularensis*.

In addition to the chitinases, we found that the KatG catalase is secreted from the LVS in a T4P-dependent manner. The function of KatG during *Francisella* infection is well understood. KatG is a catalase protein with peroxidase function, making it essential for overcoming host defense mechanisms and thereby promoting intracellular survival and replication (40). KatG has a predicted Sec signal sequence for delivery to the periplasm, but the mechanism for its release to the extracellular medium was undefined. We previously identified KatG as a prominent constituent of OMVT (19, 41), and our results now show that KatG is secreted from *Francisella* via the T4P system, likely both in soluble form to the extracellular medium and also as part of OMVT. Distinct from ChiA and ChiB, secretion of KatG was also diminished in the absence of the PilE4 putative major pilin subunit, although the dependence on PIlE4 was less than on the PilF ATPase. The LVS lacks the PIlE1-3 pilins and PilE4 may have acquired a new role in this strain, at least for some substrates such as KatG.

Our findings support a canonical role for the PilF ATPase in both T4P-mediated protein secretion and pilus fiber formation, in agreement with the prior work in *F. novicida* (28, 30). A surprise from our study was the observation of an equal loss in protein secretion for the LVS Δ*pilT* mutant, suggesting that both the PilF and PilT ATPases are required for secretion by the *F. tularensis* T4P system. In contrast to our results, loss of PilT in *F. novicida* had only a minor impact on protein secretion (30). PilT, which normally functions in fiber retraction, plays a non-canonical role in pilus assembly in *Francisella,* as T4P fiber formation is lost in both *F. novicida* and *F. tularensis* when PilT is deleted and pilus assembly is restored when the gene is complemented back (27, 28). However, we were unable to complement the secretion defects of the LVS Δ*pilT* mutant using *pilT* from either the LVS or *F. novicida*, and thus the function of PilT in *F. tularensis* secretion remains to be defined.

This study confirms that the T4P machinery functions as a hybrid pilus assembly and protein secretion machinery in *F. tularensis*. This dual function makes sense given that *Francisella* lacks secretion systems found in other intracellular pathogens and would need to maximize its secretion capabilities. We validated the secretion of ChiA, ChiD, and KatG by the LVS T4P system. The role of KatG in *F. tularensis* virulence is well documented, but understanding the functions of the different chitinases expressed by *F. tularensis* during mammalian infection may reveal new insights into the pathogenesis of tularemia. Characterization of these and other proteins secreted by the T4P system may facilitate the identification of effector proteins that underly the extreme infectivity and virulence of *F. tularensis* and its ability to evade and subvert host immune responses to cause lethal disease.

## MATERIALS AND METHODS

### Bacteria and growth conditions

All experimental studies were performed using the *Francisella tularensis* subsp. *holarctica* Live Vaccine Strain (LVS). The LVS was grown on chocolate II agar plates (BD Biosciences), in modified Mueller-Hinton Broth [MHB; Mueller-Hinton Broth (BD Biosciences) containing 1% glucose, 0.025% ferric pyrophosphate, and 0.05% L-cysteine hydrochloride](62) for plasmid transformations, or in Brain Heart Infusion (BHI) broth [37 g/L BHI powder (BD Biosciences), adjusted to pH 6.8] for all other experiments. The LVS was grown at 37°C, 5% CO2, with shaking at 100 rpm. For the BONCAT studies, bacteria were grown in the presence of 1 mM azidonorleucine (ANL). ANL was synthesized by Chem-Master International, Inc. (Hauppauge, NY). For plasmid selection, 1.25 mg/ml kanamycin or 140 µg/ml hygromycin B was added to the LVS cultures. The *Escherichia coli* DH5α strain was used for plasmid construction. DH5α was grown using Luria broth or agar, with addition of 1.25 mg/ml kanamycin or 200 µg/ml hygromycin B for plasmid selection.

### Strain and plasmid construction

Table S2 lists the bacterial strains and plasmids used in this study. Table S3 lists the primers used in this study. All plasmid constructs were verified by PCR and sequencing. All LVS strain constructs were verified by PCR.

The LVS Δ*pilE4*, Δ*pilF*, and Δ*pilT* deletion mutants were constructed by allelic exchange following the protocol described in LoVullo *et al*. (62). Briefly, ∼500 nucleotide upstream and downstream regions of each gene, separated by a 15 base pair GC-rich linker region, were cloned into the multicloning site of suicide vector pMP812 and introduced into the LVS by electroporation (62). Kanamycin resistant colonies were counter selected for sucrose resistance on MHB agar plates containing 5% sucrose. Sucrose-resistant, kanamycin-sensitive colonies were confirmed to have complete deletions of the corresponding gene by PCR.

For BONCAT analysis, the NLL-MetRS mutant methionyl-tRNA synthetase (35) was codon optimized for expression in *Francisella* and cloned into plasmid pMP831 (63), using the company GenScript (Piscataway, NJ). The promoter driving MetRS-NLL expression was then replaced with the *Francisella* RpsL promoter, generating plasmid pRpsL-MetRS_NLL. The MetRS_NLL construct was then inserted into the *att* Tn7 transposon insertion site in the chromosome of the WT LVS and Δ*pilE4*, Δ*pilF*, and Δ*pilT* deletion mutant strains, as described in LoVullo *et al*. (62).

To generate 3xFLAG-tagged versions of candidate T4P-secreted proteins, genes of interest, including their natural promoters, were amplified by PCR from LVS DNA. The PCR reaction was used to add a C-terminal 3xFLAG tag and the product was then cloned into plasmid pMP749 for insertion into the chromosomal *att* site of the WT LVS and Δ*pilE4*, Δ*pilF*, and Δ*pilT* deletion mutant strains, as described (62). Complementation plasmids for the LVS Δ*pilE4*, Δ*pilF*, and Δ*pilT* deletion mutant strains were generated by Gibson assembly (64) using the pMP831 or pMP822 vectors, for expression under native or *blaB* promoters, respectively (63). The WT *pilE4*, *pilF*, and *pilT* genes were PCR amplified from LVS DNA and inserted into SmaI- and PstI-digested vectors. The appropriate complementing or empty vector control plasmid was then transformed into the desired LVS strain by electroporation (63).

### Generation of whole bacterial lysates and cell-free supernatant fractions

To analyze protein secretion by the LVS into the extracellular medium, bacteria were cultured overnight (16-18 h) and an aliquot of the overnight culture was used to inoculate a day culture at OD_600_ = 0.06. For BONCAT experiments, ANL was added at the beginning of the day culture incubation to 1 mM final concentration. At OD_600_ = 0.3, bacteria (7 ml, approximately 9 x 10^9^ CFU) were then washed and resuspended in Opti-MEM Reduced Serum Medium (Gibco) and incubated for an additional 4 h at 37°C and 5% CO_2_, without shaking. Actual CFU were determined at the end of the 4 h incubation to ensure equal numbers of bacteria were used for the subsequent analyses. For BONCAT experiments, ANL (1 mM) was added to the Opti-MEM. At the end of the 4 h incubation, whole bacteria were collected by centrifugation (10 min, 10,000 x g, 4°C), and the supernatant fraction was removed and filtered through a 0.22 μm syringe filter (Millipore) to generate the cell-free supernatant fraction. In some experiments, OMVT were removed from the supernatant fraction by centrifugation (1 h, 100,000 x g, 4°C) and the resulting supernatant fraction was then moved to a fresh tube. For analysis of FLAG-tagged proteins, whole bacterial pellets were resuspended in 1x SDS sample buffer and heated at 95°C for 10 min. For BONCAT analysis, the whole bacterial pellets were resuspended in 1 ml PBS with 0.1% SDS and 1x Complete Protease Inhibitor Cocktail minus EDTA (Roche) and sonicated on ice for 2 min.

Remaining debris was removed by centrifugation (15 min, 21,000 x g, 4°C), and then the supernatant was moved to a fresh tube. For the cell-free supernatant fractions, protein was precipitated with 10% trichloroacetic acid for 2 h on ice, collected by centrifugation (10 min, 15,000 x g, 4°C), and the pellet was washed twice with acetone and allowed to air dry. For analysis of FLAG-tagged proteins, the dried protein pellets were resuspended in 100 µl 1x SDS sample buffer and heated at 95°C for 10 min. For BONCAT analysis, the dried protein pellets were resuspended in PBS + 2% SDS and then slowly brought to a final volume of 1 ml using PBS plus EDTA-free protease inhibitor. For BONCAT analysis, protein concentrations of both the whole bacterial lysates and cell-free supernatant fractions were determined by BCA assay (Pierce) and the samples were normalized to 0.25 µg/µl.

### Click reactions for biotin labeling of whole bacterial lysates and cell-free supernatant fractions

For BONCAT analysis, ANL-labeled proteins in the whole bacterial lysates and cell free supernatant fractions were labeled with biotin using click chemistry. Click reactions were performed by adding the following to 450 µl aliquots of the normalized lysates (0.25 µg/µl): 100 µM biotin-alkyne (Sigma), 0.1 mM CuSO_4_, 0.5 mM THTPA [Tris(3-hydroxypropyltriazolylmethyl)amine (Sigma)], 5 mM aminoguanidine, and 5 mM Na+ ascorbate. The reactions were incubated for 1.5 h at room temperature (RT) with mixing by rotation.

Reactions were quenched by addition of 10 µM EDTA.

### Gel electrophoresis and detection

For analysis of the BONCAT whole bacterial lysates and cell free supernatant fractions, 5 µg of protein was heated at 95°C in SDS sample buffer and separated by SDS-PAGE using 10% gels. For analysis of FLAG-tagged proteins, 5 µl of the samples were used for SDS-PAGE. Total protein in the gels was detected by Coomassie blue staining or silver staining using the Pierce kit for mass spectrometry, according to the manufacturer’s instructions. For blotting of ANL-labeled proteins or 3xFLAG-tagged proteins, gels were transferred to nitrocellulose membranes (Amersham) at 100 V for 1 h. Membranes were blocked in 5% bovine serum albumin in TBST (Tris-buffered saline with 0.1% Tween 20) for 1.5 h at RT. For detection of ANL-labeled proteins, membranes were incubated overnight with streptavidin- horse radish peroxidase (HRP) conjugate (Sigma) at 1:30,000 dilution. For detection of 3xFLAG-tagged proteins, membranes were incubated overnight with rabbit anti-FLAG antibodies (Sigma) at 1:2,000 dilution, and the membranes were then washed with TBST and incubated with a HRP-conjugated anti-rabbit secondary antibody (Cell Signaling) at 1:10,000 dilution for 2 h at RT. All blots were imaged by using an enhanced chemiluminescence reagent (Amersham) and ImageQuant imager (GE Life Sciences), and densitometric analysis was performed using ImageJ. Statistical calculations were performed with GraphPad Prism, using data from at least three independent experiments.

### Preparation of ANL-labeled proteins for Mass Spectrometry

For analysis of the BONCAT samples by mass spectrometry, the click reactions were precipitated with acetone for 2 h on ice, then washed once with fresh acetone to remove excess biotin. The protein pellets were then resuspended in 50 µl of 2% SDS and brought to a final volume of 500 µl with 1x PBS plus EDTA-free protease inhibitor. The samples were then added to 200 µl of streptavidin beads (GenScript), mixed by rotation for 1.5 h at RT, and the beads were then washed by inverting 4-5x with 1x PBS + 0.5% SDS followed by 3 times with 1x PBS to remove nonspecifically bound proteins. The washed beads were resuspended in a final volume of 800 µl of PBS, 600 µl of which was used for mass spectrometry analysis.

### Mass Spectrometry (MS)

Biotinylated proteins bound to the streptavidin beads were eluted with 100 µl of 5% SDS, 100 mM triethyl ammonium bicarbonate (TEAB), and 10 mM dithiothreitol (DTT) at 98°C for 20 min. The beads were removed and reduced proteins alkylated in 25 mM iodoacetamide for 30 min at RT, in the dark. Proteins were precipitated by addition of 10 μl 12% phosphoric acid, followed by 700 μl S-Trap bind/wash buffer (90% methanol, 50 mM TEAB) to produce a micro precipitate. Samples were then loaded into an S-Trap mini cartridge (Protifi), washed four times with S-Trap bind/wash buffer, and then centrifuged at 4000 x g for 1 min. The samples were digested with trypsin (20 µg) in 50 mM TEAB in a humified incubator overnight at 37°C. Peptides were eluted by sequential addition of 80 µl 50 mM TEAB, 0.2% formic acid and then 50% acetonitrile, 0.2% formic acid, each followed by centrifugation at 4000 x g for 1 min. The samples were then dried by SpeedVac and resuspended in 0.2% formic acid and peptides analyzed by C18 reverse phase LC-MS/MS. The C18 HPLC columns were prepared using a P-2000 CO_2_ laser puller (Sutter Instruments) and silica tubing (100 µm ID x 20 cm) and were self-packed with 3 µl Reprosil resin. Peptides were separated using a flow rate of 300 nl/min and a gradient elution step changing from 0.1% formic acid to 40% acetonitrile (ACN) over 90 min, followed by a 90% ACN wash and re-equilibration steps.

Peptide identification and quantitation was performed using an orbital trap instrument (Q-Exactive HF, Thermo) followed by protein database searching. Four technical replicates per sample were analyzed. Electrospray ionization was achieved using spray voltage of ∼2.3 kV. Information-dependent MS acquisitions were made using a survey scan covering m/z 375–1400 at 60,000 resolution, followed by ‘top 20’ consecutive second product ion scans at 15,000 resolution. Automatic gain control targets for MS and MS/MS were 1 x 10^6^ and 2 x 10^5^, maximum injection times were 100 and 50 ms, and an MS/MS loop size of 20 and dynamic exclusion for 20 s was used. Mass resolution cutoffs for MS and MS/MS were 10 ppm and 0.1 Da, respectively. Data files were acquired with Xcalibur (Thermo). Peptide alignments and quantitation were performed using Proteome Discoverer v3.1 software (Thermo). Protein false discovery rate (FDR) experiments were binned at 0.01 and 0.05 FDR. Peptide and peptide spectral matches (PSM) FDR cutoffs were set to 0.01. Two missed tryptic cleavages were allowed and modifications considered included static cysteine derivatization and variable deamidation (NQ), water loss (ST), oxidation (M), and biotin. Pairwise peptide log2 ratios between samples allowed label free abundance calculations and t-test, based on the background population of peptides. The *Francisella tularensis* UniProt dataset was used for data alignment. Matched peptide-based label free quantitation and *P* values were calculated by Benjamini-Hochberg correction for FDR.

### Proteomics analysis in R Studio

The nf-core/quantums pipeline was used to process raw MS data files to identify peptides, quantify proteins, and perform a differential abundance analysis (LVS Δ*pilE4* vs WT LVS and LVS Δ*pilF* vs WT LVS). Approximately 426 proteins were quantified per biological replicate (data are representative of 3 biological replicates). The protein IDs are from the *Francisella tularensis* subsp. *holarctica* LVS genome on NCBI: https://www.ncbi.nlm.nih.gov/datasets/genome/GCF_000833335.1/.

## ACKNOWLEDGEMENTS

We thank John D. Haley and the Biological Mass Spectrometry Core at Stony Brook University for performing the mass spectrometry analysis. We thank Dave Carlson and the Genomics Core Facility at Stony Brook University for bioinformatics analysis.

This work was supported by NIAID, NIH awards R01 AI141633 (D.G.T.) and T32 AI007539 (P.T.B.). The funders had no role in study design, data collection, and interpretation, or the decision to submit the work for publication.

## Notes

### Competing Interest Statement

The authors have declared no competing interest.

### Summary of Updates

This revision is to add the supplementary material or the manuscript.

