## Supplementary information for "Protein secretion by the type IV pilus machinery in *Francisella tularensis*"

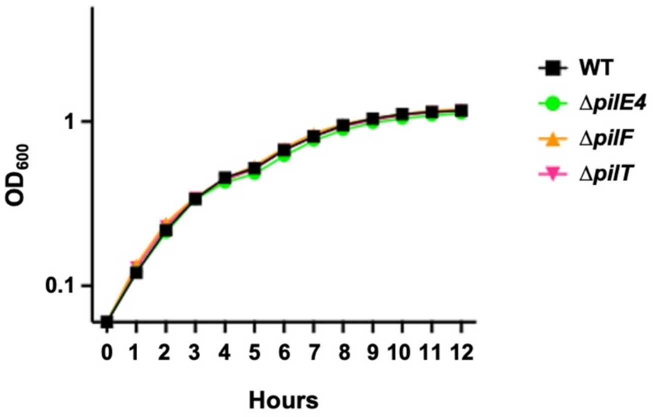


**Figure S1. Growth comparison of the LVS WT and *pil* deletion mutants.** Growth of the LVS WT and Δ*pilE4,* Δ*pilF,* and Δ*pilT* mutants in BHI medium was monitored over 12 h by recording the optical density at 600 nm ($OD_{600}$) every hour. The data are representative of two independent replicates.

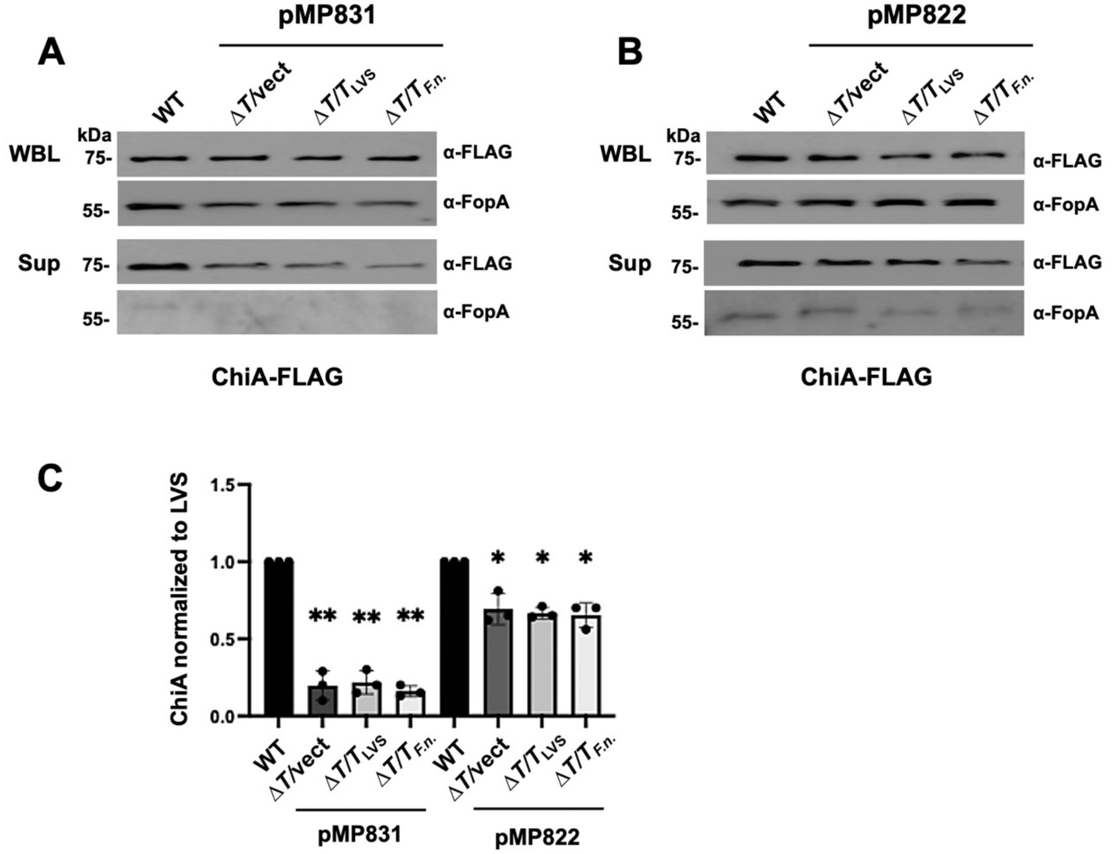


**Figure S2. Analysis of the contribution of OMVT or background lysis to ChiA secretion.** Whole bacterial lysates (WBL) and cell-free supernatant fractions (Sup) were collected from the LVS WT and Δ*pilF* (Δ*F*) and Δ*pilT* (Δ*T*) mutant strains, each expressing FLAG-tagged ChiA, following incubation in minimal infection medium. The Sup fraction was divided into two aliquots, and one aliquot was further subjected to ultracentrifugation to remove any vesicles present (Sup -OMVT). The samples were analyzed by SDS-PAGE and blotting with anti-FLAG antibodies to detect ChiA **(A)** or anti-FopA antibodies as a measure for background bacterial lysis **(B)**. **(C)** Densitometry analysis of the ChiA bands in the supernatant fractions, normalized to the WT LVS. Data represent means ± SD from three independent experiments. No significant differences were observed between the Sup and Sup -OMVT samples; calculated by one-way ANOVA with Brown-Forsythe test and pairwise t-test.

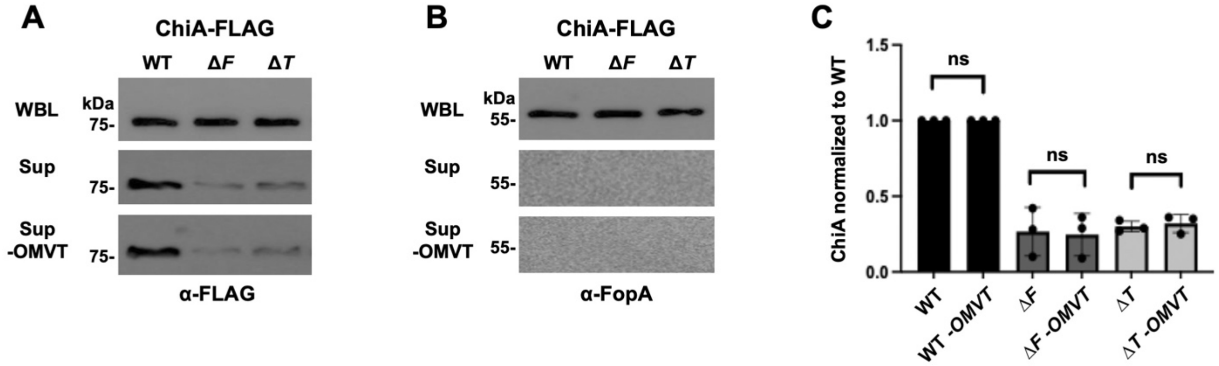


**Figure S3. Complementation of the ∆*pilT* mutant using native or over-expression promoters.** Whole bacterial lysates (WBL) and cell-free supernatant fractions (Sup) were collected from the WT LVS and ∆*pilT* (∆*T*) mutant strain, each expressing FLAG-tagged ChiA, following incubation in minimal infection medium. The mutant strain was complemented with plasmids expressing *pilT* from the LVS or *F. novicida*, or transformed with vector only as a negative control. The complementing *pilT* gene was expressed under its native promoter using plasmid pMP831 **(A),** or from a strong constitutive promoter using plasmid pMP822 **(B)**. The samples were analyzed by SDS-PAGE and blotting with anti-FLAG antibodies to detect ChiA or anti-FopA antibodies as a measure for background bacterial lysis. **(C)** Densitometry analysis of the ChiA bands in the corresponding supernatant fractions, normalized to the WT LVS. Data represent means ± SD from three independent experiments. *, $P < 0.05$; **, $P < 0.01$; calculated by one-way ANOVA with Brown-Forsythe test and pairwise t-test.

**Table S1.** LVS secreted proteins identified by BONCAT.

| Average Abundance | Locus Tag | Gene Symbol | MW [kDa] | Pfam IDs |
|---|---|---|---|---|
| 2.58E+09 | FTL_0094 | clpB | 96 | Pf00004, Pf02861, Pf07724, Pf10431, Pf17871 |
| 1.09E+09 | FTL_0519 | minD | 30.1 | Pf13614 |
| 8.73E+08 | FTL_1041 | | 36.4 | Pf00348 |
| 4.58E+08 | FTL_1714 | groEL; groL | 57.4 | Pf00118 |
| 4.52E+08 | FTL_1751 | tuf | 43.4 | Pf00009, Pf03143, Pf03144 |
| 3.90E+08 | FTL_1504 | katG | 82.5 | Pf00141 |
| 2.89E+08 | FTL_1191 | dnaK | 69.1 | Pf00012 |
| 2.67E+08 | FTL_0653 | | 15.7 | Pf02657 |
| 2.32E+08 | FTL_0009 | | 19.5 | Pf03938 |
| 1.59E+08 | FTL_0267 | htpG | 72.3 | Pf00183, Pf13589 |
| 1.03E+08 | FTL_0234 | fusA | 77.7 | Pf00009, Pf00679, Pf03144, Pf03764, Pf14492 |
| 9.29E+07 | FTL_1158 | iglB | 57.9 | Pf05943, Pf18945 |
| 8.04E+07 | FTL_1060 | | 48 | Pf00768, Pf07943 |
| 7.55E+07 | FTL_1478 | guaB | 52.1 | Pf00478, Pf00571 |
| 7.13E+07 | FTL_0546 | | 32.8 | Pf00348 |
| 7.04E+07 | FTL_0572 | | 51.9 | Pf20891 |
| 6.96E+07 | FTL_0573 | | 35.8 | Pf13036 |
| 6.84E+07 | FTL_1916 | | 76.4 | Pf03772, Pf13567 |

| | | | | |
|---|---|---|---|---|
| 6.77E+07 | FTL_1908 | ftsA | 44.8 | Pf02491, Pf14450 |
| 6.50E+07 | FTL_0269 | gdh | 49.1 | Pf00208, Pf02812 |
| 6.28E+07 | FTL_1743 | | 157.3 | |
| 6.15E+07 | FTL_0298 | | 74.2 | |
| 6.09E+07 | FTL_1592 | accB | 16.4 | Pf00364 |
| 6.04E+07 | FTL_1907 | ftsZ | 39.7 | Pf00091, Pf12327 |
| 6.02E+07 | FTL_1225 | | 25.7 | |
| 5.17E+07 | FTL_0260 | rpsD | 23.2 | Pf00163, Pf01479 |
| 5.12E+07 | FTL_1168 | | 65 | |
| 4.84E+07 | FTL_1521 | | 83.5 | |
| 4.77E+07 | FTL_1553 | sucC | 41.5 | Pf00549, Pf08442 |
| 4.53E+07 | FTL_1224 | trxA | 12.2 | Pf00085 |
| 4.48E+07 | FTL_0244 | rpmC | 7.8 | Pf00831 |
| 4.37E+07 | FTL_0309 | aceE | 100.2 | Pf00456, Pf17831 |
| 3.71E+07 | FTL_0891 | tig | 49.5 | Pf00254, Pf05697, Pf05698 |
| 3.64E+07 | FTL_0588 | | 82.3 | Pf03971 |
| 3.57E+07 | FTL_1591 | accC | 50 | Pf00289, Pf02785, Pf02786 |
| 3.40E+07 | FTL_1146 | gap | 35.4 | Pf00044, Pf02800 |
| 3.39E+07 | FTL_0617 | | 16.8 | Pf00210 |
| 3.38E+07 | FTL_1159 | iglC; iglC2 | 22.1 | Pf11550 |
| 3.33E+07 | FTL_1772 | acn; acnA | 102.6 | Pf00330, Pf00694 |
| 3.28E+07 | FTL_0225 | tsf | 30.9 | Pf00889 |
| 3.18E+07 | FTL_1912 | rpsA | 61.6 | Pf00575 |
| 3.00E+07 | FTL_0387 | | 44.4 | Pf00155 |

| | | | | |
|---|---|---|---|---|
| 2.99E+07 | FTL_0987 | mdh | 34.1 | Pf00056, Pf02866 |
| 2.87E+07 | FTL_1139 | fabG | 26.3 | Pf13561 |
| 2.63E+07 | FTL_0020 | | 67 | |
| 2.46E+07 | FTL_1789 | gltA | 46.8 | Pf00285 |
| 2.45E+07 | FTL_1461 | deoD | 26.8 | Pf01048 |
| 2.43E+07 | FTL_1744 | rpoB | 151.2 | Pf00562, Pf04560, Pf04561, Pf04563, Pf04565, Pf10385 |
| 2.39E+07 | FTL_1389 | nadC | 31.5 | Pf01729, Pf02749 |
| 2.35E+07 | FTL_1597 | lptD | 98.5 | Pf03968, Pf04453 |
| 2.34E+07 | FTL_1404 | rplT | 13.3 | Pf00453 |
| 2.20E+07 | FTL_1308 | | 44.9 | Pf08245 |
| 2.19E+07 | FTL_1015 | | 19.7 | Pf08534 |
| 2.17E+07 | FTL_0789 | | 44.2 | Pf00155 |
| 1.96E+07 | FTL_1061 | ppa | 19.6 | Pf00719 |
| 1.87E+07 | FTL_1423 | | 38.9 | Pf00535 |
| 1.83E+07 | FTL_1746 | rplJ | 18.7 | Pf00466 |
| 1.81E+07 | FTL_1605 | sohB | 38 | Pf01343, Pf08496 |
| 1.81E+07 | FTL_0207 | pcp | 24.2 | Pf01470 |
| 1.74E+07 | FTL_1442 | fabI | 27.8 | Pf13561 |
| 1.73E+07 | FTL_1137 | fabF | 44 | Pf00109, Pf02801 |
| 1.67E+07 | FTL_1780 | tpiA | 27.6 | Pf00121 |
| 1.63E+07 | FTL_0536 | | 18.8 | Pf03938 |
| 1.62E+07 | FTL_1701 | glpX | 34.8 | Pf03320 |
| 1.59E+07 | FTL_0661 | | 59.9 | |
| 1.57E+07 | FTL_0477 | gcvT | 39.5 | Pf01571, Pf08669 |

| | | | | |
|---|---|---|---|---|
| 1.56E+07 | FTL_0534 | dxr | 42.8 | Pf02670, Pf08436, Pf13288 |
| 1.56E+07 | FTL_1783 | odhB | 52.7 | Pf00198, Pf00364, Pf02817 |
| 1.53E+07 | FTL_1490 | gpmI | 57.6 | Pf01676, Pf06415 |
| 1.50E+07 | FTL_1162 | | 155.8 | |
| 1.34E+07 | FTL_1715 | groES; groS | 10.3 | Pf00166 |
| 1.33E+07 | FTL_1140 | fabD | 33.5 | Pf00698 |
| 1.32E+07 | FTL_0253 | rpsE | 17.5 | Pf00333, Pf03719 |
| 1.30E+07 | FTL_1024 | rpsF | 13 | Pf01250 |
| 1.29E+07 | FTL_0227 | frr | 20.5 | Pf01765 |
| 1.26E+07 | FTL_0862 | selO | 52.5 | Pf02696 |
| 1.24E+07 | FTL_0596 | | 48.8 | Pf00984, Pf03720, Pf03721 |
| 1.23E+07 | FTL_1522 | kbl | 43.9 | Pf00155 |
| 1.21E+07 | FTL_0569 | omp26 | 19.8 | |
| 1.20E+07 | FTL_0533 | gyrA | 97 | Pf00521, Pf03989 |
| 1.16E+07 | FTL_1745 | rplL | 12.8 | Pf00542, Pf16320 |
| 1.11E+07 | FTL_1148 | pyk | 51.7 | Pf00224, Pf02887 |
| 1.09E+07 | FTL_1042 | | 29.3 | Pf00254, Pf01346 |
| 1.08E+07 | FTL_1784 | | 105.6 | |
| 1.05E+07 | FTL_1407 | thrS | 72.3 | Pf00587, Pf02824, Pf03129, Pf07973 |
| 1.05E+07 | FTL_1157 | tssB | 20.9 | Pf05591 |
| 9.58E+06 | FTL_0248 | rplE | 20 | Pf00281, Pf00673 |

| | | | | |
|---|---|---|---|---|
| 9.24E+06 | FTL_0428 | parB | 34.6 | Pf02195, Pf08535 |
| 9.16E+06 | FTL_0357 | rph | 25.4 | Pf01138, Pf03725 |
| 9.06E+06 | FTL_0002 | dnaN | 41.6 | Pf00712, Pf02767, Pf02768 |
| 8.88E+06 | FTL_1546 | pdxS | 30.8 | Pf01680 |
| 8.86E+06 | FTL_0334 | tolB | 48.4 | Pf07676 |
| 8.83E+06 | FTL_0311 | lpdA | 50.5 | Pf02852, Pf07992 |
| 8.82E+06 | FTL_1554 | sucD | 30.1 | Pf00549, Pf02629 |
| 8.82E+06 | FTL_1786 | sdhA | 65.8 | Pf00890, Pf02910 |
| 8.74E+06 | FTL_0242 | rpsC | 24.9 | Pf00189, Pf07650 |
| 8.73E+06 | FTL_0766 | ggt | 65.1 | Pf01019 |
| 8.63E+06 | FTL_0433 | rlmE | 23.2 | Pf01728 |
| 8.57E+06 | FTL_1197 | pheS | 38.5 | Pf01409, Pf02912 |
| 8.41E+06 | FTL_1527 | eno | 49.5 | Pf00113, Pf03952 |
| 8.29E+06 | FTL_0224 | rpsB | 26.4 | Pf00318 |
| 8.24E+06 | FTL_0295 | accA | 35.4 | Pf03255 |
| 8.04E+06 | FTL_1678 | | 38.4 | Pf02016, Pf17676 |
| 7.64E+06 | FTL_1672 | | 112.4 | Pf00873 |
| 7.53E+06 | FTL_1547 | gyrB | 89.7 | Pf00204, Pf00986, Pf01751, Pf02518, Pf18053, Pf21249 |
| 7.53E+06 | FTL_1747 | rplA | 24.5 | Pf00687 |
| 7.49E+06 | FTL_1748 | rplK | 15.3 | Pf00298, Pf03946 |
| 7.41E+06 | FTL_0070 | | 10.1 | |

| | | | | |
|---|---|---|---|---|
| 7.36E+06 | FTL_1492 | | 32.8 | Pf00480 |
| 7.29E+06 | FTL_0703 | glyA | 45.3 | Pf00464 |
| 7.17E+06 | FTL_1899 | | 38.2 | Pf00120, Pf03951 |
| 7.01E+06 | FTL_0310 | aceF | 56.8 | Pf00198, Pf00364, Pf02817 |
| 6.89E+06 | FTL_1598 | argS | 65.9 | Pf00750, Pf03485, Pf05746 |
| 6.83E+06 | FTL_0210 | valS | 104.7 | Pf00133, Pf08264, Pf10458 |
| 6.82E+06 | FTL_1479 | pepA | 52 | Pf00883, Pf02789 |
| 6.64E+06 | FTL_0028 | pyrB | 35.1 | Pf00185, Pf02729 |
| 6.62E+06 | FTL_0968 | tyrS | 44.6 | Pf00579, Pf01479 |
| 6.60E+06 | FTL_0538 | fabZ | 18.1 | Pf07977 |
| 6.57E+06 | FTL_1476 | pgi | 61.1 | Pf00342 |
| 6.49E+06 | FTL_1213 | | 73.3 | Pf01841, Pf12969 |
| 6.14E+06 | FTL_0480 | | 52.7 | Pf02347, Pf21478 |
| 6.02E+06 | FTL_0610 | rho | 47.1 | Pf00006, Pf07497, Pf07498 |
| 5.97E+06 | FTL_1248 | gor; gorA | 49.5 | Pf02852, Pf07992 |
| 5.95E+06 | FTL_0261 | | 35.3 | |
| 5.67E+06 | FTL_1283 | hemL | 47 | Pf00202 |
| 5.66E+06 | FTL_1306 | | 39.9 | Pf08238 |
| 5.55E+06 | FTL_1392 | deaD | 64 | Pf00270, Pf00271, Pf03880 |
| 5.20E+06 | FTL_0259 | rpsK | 13.8 | Pf00411 |

| | | | | |
|---|---|---|---|---|
| 5.18E+06 | FTL_0583 | fadA | 41.7 | Pf00108, Pf02803 |
| 5.14E+06 | FTL_1968 | | 57.5 | Pf00575, Pf10150, Pf20833 |
| 5.04E+06 | FTL_0986 | | 66.5 | Pf00317, Pf02867 |
| 5.03E+06 | FTL_1309 | accD | 33.4 | Pf01039, Pf17848 |
| 4.98E+06 | FTL_1579 | | 25.7 | |
| 4.92E+06 | FTL_1142 | plsX | 37.8 | Pf02504 |
| 4.86E+06 | FTL_0597 | | 36.4 | |
| 4.82E+06 | FTL_0237 | | 22.5 | |
| 4.80E+06 | FTL_0462 | parC | 83.9 | Pf00521, Pf03989 |
| 4.76E+06 | FTL_0895 | hupB | 9.5 | Pf00216 |
| 4.70E+06 | FTL_1310 | ndk | 15.4 | Pf00334 |
| 4.65E+06 | FTL_1832 | fslA | 74.1 | Pf04183, Pf06276 |
| 4.58E+06 | FTL_1025 | rpsR | 8.4 | Pf01084 |
| 4.51E+06 | FTL_0250 | rpsH | 14.4 | Pf00410 |
| 4.47E+06 | FTL_0984 | | 47.3 | Pf00268, Pf00462 |
| 4.44E+06 | FTL_0584 | | 100.6 | Pf00378, Pf00725, Pf00887, Pf02737 |
| 4.38E+06 | FTL_0776 | | 29.8 | |
| 4.16E+06 | FTL_1147 | pgk | 42 | Pf00162 |
| 4.16E+06 | FTL_1943 | priA | 81 | Pf00270, Pf17764, Pf18074 |
| 4.09E+06 | FTL_1190 | grpE | 22 | Pf01025 |
| 4.08E+06 | FTL_0890 | | 22.4 | |
| 4.07E+06 | FTL_0493 | | 29.6 | |

| | | | | |
|---|---|---|---|---|
| 4.04E+06 | FTL_0072 | mnmA; trmU | 40.1 | Pf03054, Pf20258, Pf20259 |
| 3.98E+06 | FTL_1179 | raiA | 11.1 | Pf02482 |
| 3.87E+06 | FTL_1212 | leuS | 93.3 | Pf00133, Pf08264, Pf09334, Pf13603 |
| 3.87E+06 | FTL_1958 | trpCF | 51.3 | Pf00218, Pf00697 |
| 3.84E+06 | FTL_0964 | hslU | 51.2 | Pf00004, Pf07724 |
| 3.75E+06 | FTL_0525 | | 55 | Pf05681, Pf05683 |
| 3.72E+06 | FTL_1791 | sodB | 21.9 | Pf00081, Pf02777 |
| 3.70E+06 | FTL_0730 | | 29.8 | Pf08282 |
| 3.68E+06 | FTL_1812 | hemE | 38.8 | Pf01208 |
| 3.64E+06 | FTL_0001 | dnaA | 55.8 | Pf00308, Pf08299, Pf11638 |
| 3.62E+06 | FTL_1141 | | 35.2 | |
| 3.61E+06 | FTL_1474 | greA | 17.7 | Pf01272, Pf03449 |
| 3.56E+06 | FTL_0894 | lon | 86.2 | Pf00004, Pf02190, Pf05362 |
| 3.53E+06 | FTL_0044 | | 129.1 | |
| 3.50E+06 | FTL_0743 | | 26 | Pf00106 |
| 3.49E+06 | FTL_0294 | mutS | 95.8 | Pf00488, Pf01624, Pf05188, Pf05190, Pf05192 |
| 3.49E+06 | FTL_1491 | serS | 48.5 | Pf00587, Pf02403 |
| 3.46E+06 | FTL_0454 | glmS | 67.4 | Pf01380, Pf13522 |
| 3.45E+06 | FTL_1532 | | 21 | |

| | | | | |
|---|---|---|---|---|
| 3.42E+06 | FTL_1495 | cydD | 65.8 | Pf00005, Pf00664 |
| 3.33E+06 | FTL_0436 | ileS | 106.9 | Pf00133, Pf06827, Pf08264 |
| 3.32E+06 | FTL_1683 | cysS | 52.4 | Pf01406, Pf09190 |
| 3.22E+06 | FTL_1229 | sufC | 27.4 | Pf00005 |
| 3.07E+06 | FTL_1458 | secA | 103.5 | Pf01043, Pf02810, Pf07516, Pf07517, Pf21090 |
| 3.06E+06 | FTL_1272 | bioB | 34.9 | Pf04055, Pf06968 |
| 3.05E+06 | FTL_1350 | glyS | 78 | Pf02092, Pf05746 |
| 2.99E+06 | FTL_1240 | | 40.9 | Pf00793 |
| 2.97E+06 | FTL_0514 | | 25.2 | Pf05494 |
| 2.92E+06 | FTL_1860 | | 141.3 | |
| 2.86E+06 | FTL_1523 | tdh | 38.4 | Pf00107, Pf08240 |
| 2.85E+06 | FTL_0898 | hfq | 12.5 | Pf17209 |
| 2.80E+06 | FTL_0166 | | 30.2 | Pf00582 |
| 2.77E+06 | FTL_0479 | gcvPA | 49.6 | Pf02347 |
| 2.75E+06 | FTL_1276 | | 31 | Pf03099, Pf08279 |
| 2.72E+06 | FTL_1149 | fba | 38.1 | Pf01116 |
| 2.69E+06 | FTL_0029 | carB | 120.6 | Pf02142, Pf02786, Pf02787 |
| 2.64E+06 | FTL_0768 | typA | 67.6 | Pf00009, Pf00679, Pf03144, Pf21018 |
| 2.60E+06 | FTL_0171 | | 32.4 | |
| 2.57E+06 | FTL_0780 | | 16.2 | |

| | | | | |
|---|---|---|---|---|
| 2.56E+06 | FTL_1904 | prfB | 36.6 | Pf00472, Pf03462 |
| 2.54E+06 | FTL_0805 | putA | 149.9 | Pf00171, Pf01619, Pf14850, Pf18327 |
| 2.47E+06 | FTL_1266 | bioJ | 35.5 | Pf07859 |
| 2.44E+06 | FTL_1795 | atpD | 49.8 | Pf00006, Pf02874 |
| 2.41E+06 | FTL_0823 | | 13.1 | |
| 2.40E+06 | FTL_0012 | recA | 38.8 | Pf00154, Pf21096 |
| 2.35E+06 | FTL_0251 | rplF; rplF_2 | 19.1 | Pf00347 |
| 2.34E+06 | FTL_1453 | rplU | 11.6 | Pf00829 |
| 2.30E+06 | FTL_0246 | rplN | 13.2 | Pf00238 |
| 2.27E+06 | FTL_1117 | | 31.1 | |
| 2.21E+06 | FTL_0739 | gidA; mnmG | 69.7 | Pf01134, Pf13932, Pf21680 |
| 2.21E+06 | FTL_1145 | tkt | 73.3 | Pf00456, Pf02779, Pf02780 |
| 2.20E+06 | FTL_1809 | infB | 92.4 | Pf00009, Pf04760, Pf11987 |
| 2.18E+06 | FTL_1230 | sufB | 53.5 | Pf01458, Pf19295 |
| 2.17E+06 | FTL_1726 | parE | 70.3 | Pf00204, Pf00986, Pf01751, Pf02518 |
| 2.16E+06 | FTL_0886 | miaB | 50.1 | Pf00919, Pf01938, Pf04055 |
| 2.11E+06 | FTL_1850 | purB | 49.4 | Pf00206 |
| 2.11E+06 | FTL_1334 | sdaA | 50 | Pf03313, Pf03315 |
| 2.10E+06 | FTL_0540 | lpxB | 43.1 | Pf02684 |

| | | | | |
|---|---|---|---|---|
| 2.09E+06 | FTL_1026 | rplI | 16.1 | Pf01281, Pf03948 |
| 2.08E+06 | FTL_0674 | panB | 28.9 | Pf02548 |
| 2.07E+06 | FTL_0483 | glgB | 74.9 | Pf00128, Pf02806, Pf02922 |
| 2.07E+06 | FTL_1571 | trxB | 34 | Pf07992 |
| 2.00E+06 | FTL_0438 | | 67.3 | Pf00390, Pf03949 |
| 1.87E+06 | FTL_0717 | rne | 95.9 | Pf00575, Pf10150, Pf20833 |
| 1.84E+06 | FTL_0616 | rpoA; rpoA2 | 35.1 | Pf01000, Pf01193, Pf03118 |
| 1.83E+06 | FTL_0236 | rplC | 22.3 | Pf00297 |
| 1.82E+06 | FTL_1135 | mnmD | 26.7 | Pf05430 |
| 1.80E+06 | FTL_1910 | ddl | 32.7 | Pf07478 |
| 1.75E+06 | FTL_1874 | cgtA; obg | 36.9 | Pf01018, Pf01926 |
| 1.71E+06 | FTL_0333 | tolA | 34.6 | |
| 1.70E+06 | FTL_0419 | | 76.5 | Pf01432, Pf19310 |
| 1.69E+06 | FTL_0552 | | 25.5 | Pf00072, Pf00486 |
| 1.68E+06 | FTL_1049 | dnaG | 69 | Pf01807, Pf08275, Pf10410, Pf13155 |
| 1.65E+06 | FTL_0994 | | 56 | |
| 1.62E+06 | FTL_1903 | | 66.2 | |
| 1.62E+06 | FTL_1494 | | 18.1 | |
| 1.61E+06 | FTL_0655 | | 40 | |
| 1.58E+06 | FTL_0522 | rpmB | 8.9 | Pf00830 |
| 1.58E+06 | FTL_1582 | | 45.2 | Pf13416 |
| 1.57E+06 | FTL_1050 | rpoD | 67.6 | Pf00140, Pf03979, Pf04539, |

| | | | | |
|---|---|---|---|---|
| | | | | Pf04542, Pf04545, Pf04546 |
| 1.55E+06 | FTL_1738 | rpsP | 9.1 | Pf00886 |
| 1.52E+06 | FTL_1411 | | 12.3 | Pf02575 |
| 1.49E+06 | FTL_1537 | | 75.4 | |
| 1.47E+06 | FTL_0762 | | 16.5 | Pf03364 |
| 1.43E+06 | FTL_1646 | | 22.7 | |
| 1.41E+06 | FTL_0795 | adk | 24.4 | Pf00406, Pf05191 |
| 1.39E+06 | FTL_0928 | elbB | 23.7 | Pf01965 |
| 1.36E+06 | FTL_0605 | rfbA | 32.4 | Pf00483 |
| 1.34E+06 | FTL_1966 | trpE | 58 | Pf00425, Pf04715 |
| 1.33E+06 | FTL_0238 | rplW | 11.1 | Pf00276 |
| 1.33E+06 | FTL_0174 | | 16.7 | |
| 1.31E+06 | FTL_0831 | cphA | 103.9 | Pf02786, Pf02875, Pf08245, Pf18921 |
| 1.31E+06 | FTL_1892 | | 39.2 | Pf04488 |
| 1.30E+06 | FTL_1896 | | 50.9 | |
| 1.30E+06 | FTL_0875 | | 43.9 | |
| 1.29E+06 | FTL_0281 | | 33.6 | Pf00226, Pf01556 |
| 1.24E+06 | FTL_0834 | | 27.8 | Pf00581 |
| 1.21E+06 | FTL_0254 | rpmD | 6.9 | Pf00327 |
| 1.20E+06 | FTL_1393 | ppiC | 10.2 | Pf00639 |
| 1.18E+06 | FTL_1671 | | 50 | |
| 1.18E+06 | FTL_0650 | proS | 63.4 | Pf00587, Pf03129, Pf04073 |
| 1.17E+06 | FTL_1109 | | 38 | |
| 1.17E+06 | FTL_0277 | | 51.5 | Pf02852, Pf07992 |

| | | | | |
|---|---|---|---|---|
| 1.16E+06 | FTL_1438 | | 11.2 | |
| 1.16E+06 | FTL_0343 | | 59.3 | Pf04244 |
| 1.15E+06 | FTL_0601 | wbtI | 40.7 | Pf01041 |
| 1.15E+06 | FTL_1810 | nusA | 55.1 | Pf00575, Pf08529, Pf13184, Pf14520 |
| 1.14E+06 | FTL_1956 | pepN | 98 | Pf01433, Pf11940, Pf17432, Pf17900 |
| 1.12E+06 | FTL_1108 | | 51.3 | Pf00883, Pf21337 |
| 1.08E+06 | FTL_0916 | ilvC | 37.9 | Pf01450, Pf07991 |
| 1.07E+06 | FTL_1418 | | 30.9 | Pf01112 |
| 1.05E+06 | FTL_1027 | | 52 | |
| 1.04E+06 | FTL_1468 | hflB | 70.7 | Pf00004, Pf01434, Pf06480, Pf17862 |
| 1.03E+06 | FTL_0877 | | 67.4 | |
| 1.03E+06 | FTL_0239 | rplB | 30.4 | Pf00181, Pf03947 |
| 1.03E+06 | FTL_1420 | | 40.3 | Pf00294 |
| 1.02E+06 | FTL_0892 | clpP | 22.1 | Pf00574 |
| 1.01E+06 | FTL_0832 | | 63.1 | Pf02875, Pf08245 |
| 9.90E+05 | FTL_0033 | | 50.3 | Pf01979 |
| 9.88E+05 | FTL_1185 | | 23.6 | |
| 9.87E+05 | FTL_0233 | rpsG | 17.8 | Pf00177 |
| 9.70E+05 | FTL_1311 | pyrG | 61 | Pf00117, Pf06418 |
| 9.65E+05 | FTL_1785 | | 26.5 | Pf13085, Pf13183 |
| 9.59E+05 | FTL_0218 | gltX | 53 | Pf00749, Pf19269 |

| | | | | |
|---|---|---|---|---|
| 9.50E+05 | FTL_1603 | yhbY | 10.4 | Pf01985 |
| 9.45E+05 | FTL_0489 | glyQ | 34.3 | Pf02091 |
| 8.99E+05 | FTL_1824 | | 87.3 | |
| 8.97E+05 | FTL_0258 | rpsM | 13.4 | Pf00416 |
| 8.84E+05 | FTL_1187 | rplM | 15.9 | Pf00572 |
| 8.83E+05 | FTL_0262 | rplQ | 16.8 | Pf01196 |
| 8.74E+05 | FTL_1706 | lolA | 23.4 | Pf03548 |
| 8.53E+05 | FTL_0249 | rpsN | 11.7 | Pf00253 |
| 8.48E+05 | FTL_0076 | ribB | 44.6 | Pf00925, Pf00926 |
| 8.33E+05 | FTL_0960 | sthA | 52.3 | Pf02852, Pf07992 |
| 8.23E+05 | FTL_0132 | ppdK | 97.6 | Pf00391, Pf01326, Pf02896 |
| 8.21E+05 | FTL_0074 | def | 19.8 | Pf01327 |
| 8.21E+05 | FTL_0473 | def | 24.2 | Pf01327 |
| 8.14E+05 | FTL_0487 | glgP | 86.4 | Pf00343 |
| 8.00E+05 | FTL_1198 | pheT | 88.1 | Pf01588, Pf03147, Pf03483, Pf03484, Pf17759 |
| 7.99E+05 | FTL_1842 | gatA | 52.4 | Pf01425 |
| 7.91E+05 | FTL_0949 | prs; prsA | 34.9 | Pf13793, Pf14572 |
| 7.81E+05 | FTL_1617 | glnS | 63.6 | Pf00749, Pf03950, Pf20974 |
| 7.79E+05 | FTL_1071 | guaA | 57.7 | Pf00117, Pf00958, Pf02540 |
| 7.74E+05 | FTL_0395 | purM | 37.7 | Pf00586, Pf02769 |
| 7.73E+05 | FTL_1189 | | 44.8 | |
| 7.71E+05 | FTL_1511 | | 38.8 | Pf03009 |
| 7.59E+05 | FTL_1419 | cphB | 29.3 | Pf03575 |

| | | | | |
|---|---|---|---|---|
| 7.45E+05 | FTL_0485 | glgC | 47.5 | Pf00483 |
| 7.41E+05 | FTL_1793 | chiD | 105.3 | Pf00704 |
| 7.39E+05 | FTL_1841 | | 53 | |
| 7.37E+05 | FTL_0014 | ssb | 17.5 | Pf00436 |
| 7.33E+05 | FTL_0893 | clpX | 46.3 | Pf06689, Pf07724, Pf10431 |
| 7.31E+05 | FTL_1906 | lpxC | 31.8 | Pf03331 |
| 7.17E+05 | FTL_1302 | | 27.7 | Pf00175, Pf00970 |
| 7.13E+05 | FTL_1861 | purF | 55.4 | Pf00156, Pf13522 |
| 7.11E+05 | FTL_1930 | purA | 46.9 | Pf00709 |
| 7.08E+05 | FTL_1648 | | 29.5 | Pf00005, Pf08352 |
| 7.03E+05 | FTL_1781 | glmM | 48.2 | Pf00408, Pf02878, Pf02879, Pf02880 |
| 7.02E+05 | FTL_1129 | | 18.9 | |
| 7.02E+05 | FTL_0549 | proC | 30.1 | Pf03807, Pf14748 |
| 6.88E+05 | FTL_1239 | ffh | 50.3 | Pf00448, Pf02881, Pf02978 |
| 6.84E+05 | FTL_0852 | aroA | 46.8 | Pf00275 |
| 6.78E+05 | FTL_1482 | | 47.8 | Pf00675, Pf05193 |
| 6.65E+05 | FTL_1929 | purH | 56.3 | Pf01808, Pf02142 |
| 6.46E+05 | FTL_0187 | | 27.9 | Pf00497 |
| 6.46E+05 | FTL_0133 | feoB | 81.4 | Pf02421, Pf07664, Pf07670 |
| 6.31E+05 | FTL_1328 | | 41.2 | Pf00691 |
| 6.29E+05 | FTL_0923 | grxB | 25.1 | Pf04399, Pf13417 |

| | | | | |
|---|---|---|---|---|
| 6.20E+05 | FTL_1628 | | 25.9 | |
| 6.03E+05 | FTL_1733 | | 32.3 | Pf03171 |
| 6.02E+05 | FTL_1498 | | 13.7 | Pf01042 |
| 6.01E+05 | FTL_1524 | | 47.7 | Pf03222 |
| 5.96E+05 | FTL_1473 | uvrA | 105 | Pf17755, Pf17760 |
| 5.87E+05 | FTL_1186 | rpsI | 14.7 | Pf00380 |
| 5.77E+05 | FTL_0404 | glk | 37.5 | Pf02685 |
| 5.71E+05 | FTL_0240 | | 10.5 | |
| 5.66E+05 | FTL_1166 | | 55.3 | |
| 5.65E+05 | FTL_1107 | | 52.4 | |
| 5.37E+05 | FTL_0537 | lpxD; lpxD2 | 35.4 | Pf00132, Pf04613, Pf14602 |
| 5.36E+05 | FTL_0521 | rpmG | 6.1 | Pf00471 |
| 5.18E+05 | FTL_1160 | | 46.4 | |
| 5.16E+05 | FTL_1602 | hemB | 35.8 | Pf00490 |
| 5.02E+05 | FTL_1228 | sufD | 43.2 | Pf01458 |
| 5.02E+05 | FTL_1304 | gshA | 56.9 | Pf04262 |
| 4.96E+05 | FTL_0801 | aroK | 19.7 | Pf01202 |
| 4.96E+05 | FTL_0927 | lipA | 36.8 | Pf04055, Pf16881 |
| 4.93E+05 | FTL_0255 | rplO | 15.1 | Pf00828 |
| 4.85E+05 | FTL_1900 | glsA | 57.2 | Pf04960, Pf12796, Pf17959 |
| 4.84E+05 | FTL_0306 | trpS | 37.9 | Pf00579 |
| 4.79E+05 | FTL_0938 | | 43.4 | Pf00282 |
| 4.72E+05 | FTL_0611 | trxA | 12 | Pf00085 |
| 4.64E+05 | FTL_1018 | serC | 39.3 | Pf00266 |
| 4.40E+05 | FTL_0933 | rmuC | 54.5 | Pf02646 |
| 4.35E+05 | FTL_0421 | | 15.8 | |
| 4.30E+05 | FTL_1132 | suhB | 28.9 | Pf00459 |
| 4.27E+05 | FTL_0232 | rpsL | 13.8 | Pf00164 |

| | | | | |
|---|---|---|---|---|
| 4.19E+05 | FTL_0034 | | 40.5 | Pf14213 |
| 4.18E+05 | FTL_1834 | | 47.4 | Pf00278, Pf02784 |
| 4.04E+05 | FTL_1390 | nadA | 38 | Pf02445 |
| 3.90E+05 | FTL_0732 | gloA | 14.4 | Pf00903 |
| 3.89E+05 | FTL_0399 | purK | 40.3 | Pf02222, Pf17769 |
| 3.85E+05 | FTL_0223 | | 21.5 | Pf00186 |
| 3.83E+05 | FTL_0926 | | 19.1 | Pf00210 |
| 3.62E+05 | FTL_0182 | efp | 20.9 | Pf01132, Pf08207, Pf09285 |
| 3.55E+05 | FTL_0466 | | 76.9 | |
| 3.43E+05 | FTL_1048 | | 16.8 | Pf09424 |
| 3.42E+05 | FTL_0767 | | 29.3 | Pf08241 |
| 3.40E+05 | FTL_1138 | acpP | 10.7 | Pf00550 |
| 3.37E+05 | FTL_0045 | | 23.9 | |
| 3.35E+05 | FTL_1245 | | 25.2 | Pf13561 |
| 3.31E+05 | FTL_0520 | minC | 24.9 | Pf03775, Pf05209 |
| 3.28E+05 | FTL_1262 | pabB | 68.3 | Pf00425, Pf01063 |
| 3.22E+05 | FTL_0673 | panC | 29.7 | Pf02569 |
| 3.21E+05 | FTL_0459 | map | 28.4 | Pf00557 |
| 3.21E+05 | FTL_1721 | prfA | 40.4 | Pf00472, Pf03462 |
| 3.16E+05 | FTL_0235 | rpsJ | 11.9 | Pf00338 |
| 3.15E+05 | FTL_0884 | ybeY | 18.6 | Pf02130 |
| 3.11E+05 | FTL_0846 | | 21.3 | Pf00857 |
| 3.11E+05 | FTL_0484 | | 59.7 | Pf02878, Pf02879, Pf02880 |
| 3.08E+05 | FTL_0098 | trpA | 29.1 | Pf00290 |
| 3.06E+05 | FTL_1275 | bioD | 24.5 | Pf13500 |
| 2.92E+05 | FTL_1664 | deoB | 45.7 | Pf01676 |

| | | | | |
|---|---|---|---|---|
| 2.90E+05 | FTL_1669 | | 41.4 | Pf01743, Pf12627 |
| 2.90E+05 | FTL_0377 | aroC | 38 | Pf01264 |
| 2.89E+05 | FTL_0929 | | 26.9 | Pf01709, Pf20772 |
| 2.83E+05 | FTL_0131 | ilvE | 33.1 | Pf01063 |
| 2.83E+05 | FTL_1606 | | 24 | |
| 2.78E+05 | FTL_0478 | gcvH | 13.9 | Pf01597 |
| 2.67E+05 | FTL_0856 | | 25.4 | Pf00484 |
| 2.64E+05 | FTL_1866 | pcm | 23.2 | Pf01135 |
| 2.60E+05 | FTL_0396 | purD | 86.5 | Pf01071, Pf01259, Pf02843, Pf02844 |
| 2.50E+05 | FTL_1093 | msrA | 32.6 | Pf01625, Pf01641 |
| 2.48E+05 | FTL_0804 | | 11.3 | Pf03937 |
| 2.48E+05 | FTL_1256 | | 16.4 | |
| 2.46E+05 | FTL_1394 | | 51.2 | Pf00083 |
| 2.45E+05 | FTL_0011 | secB; secB1; secB2 | 16.9 | Pf02556 |
| 2.34E+05 | FTL_0327 | | 35.1 | |
| 2.31E+05 | FTL_0850 | | 18.8 | |
| 2.31E+05 | FTL_0031 | | 39.5 | Pf00328 |
| 2.24E+05 | FTL_1119 | | 32.9 | Pf04381 |
| 2.23E+05 | FTL_1333 | sufS | 45.1 | Pf00266 |
| 2.22E+05 | FTL_1192 | | 43.9 | |
| 2.19E+05 | FTL_0413 | | 48.5 | Pf00275 |
| 2.14E+05 | FTL_0247 | rplX | 11.5 | Pf00467, Pf17136 |
| 2.13E+05 | FTL_1265 | | 48.2 | |
| 2.04E+05 | FTL_0965 | hslV | 19.8 | Pf00227 |
| 1.89E+05 | FTL_0602 | | 27.8 | Pf00551, Pf18216 |

| | | | | |
|---|---|---|---|---|
| 1.88E+05 | FTL_0307 | coaE | 23.4 | Pf01121 |
| 1.73E+05 | FTL_1397 | galK | 43.4 | Pf00288, Pf08544, Pf10509 |
| 1.72E+05 | FTL_1797 | atpA | 55.5 | Pf00006, Pf00306, Pf02874 |
| 1.71E+05 | FTL_1164 | | 44.6 | |
| 1.70E+05 | FTL_0864 | | 37.2 | |
| 1.67E+05 | FTL_0016 | pta | 77.1 | Pf01515, Pf07085 |
| 1.62E+05 | FTL_0492 | murF | 50.1 | Pf08245 |
| 1.61E+05 | FTL_0556 | | 85.9 | |
| 1.60E+05 | FTL_0637 | | 25.6 | Pf00753 |
| 1.59E+05 | FTL_1273 | | 42.6 | Pf00155 |
| 1.56E+05 | FTL_0200 | | 35.5 | |
| 1.54E+05 | FTL_1202 | | 36.9 | |
| 1.36E+05 | FTL_0015 | ackA | 42.2 | Pf00871 |
| 1.34E+05 | FTL_0941 | | 19.1 | Pf00857 |
| 1.27E+05 | FTL_1539 | ftsI | 62.7 | Pf00905, Pf03717 |
| 1.26E+05 | FTL_0093 | | 78.9 | Pf00704 |
| 1.25E+05 | FTL_1585 | recJ | 65.6 | Pf01368, Pf02272, Pf17768 |
| 1.17E+05 | FTL_0394 | folD | 30.5 | Pf00763, Pf02882 |
| 1.09E+05 | FTL_0740 | | 42.6 | |
| 1.09E+05 | FTL_0245 | rpsQ | 9.8 | Pf00366 |
| 1.04E+05 | FTL_0879 | blaFTU | 31.9 | Pf13354 |
| 1.02E+05 | FTL_1430 | galE | 37.8 | Pf01370 |
| 9.76E+04 | FTL_1396 | galT | 39.7 | Pf01087, Pf02744 |
| 8.85E+04 | FTL_1075 | | 22.4 | |
| 8.39E+04 | FTL_0449 | | 13.3 | |

| | | | | |
|---|---|---|---|---|
| 8.01E+04 | FTL_1106 | alaS | 96 | Pf01411, Pf02272, Pf07973 |
| 6.87E+04 | FTL_0675 | panG | 27.5 | Pf10728 |
| 5.77E+04 | FTL_0802 | | 40.1 | |
| 5.10E+04 | FTL_1710 | | 27.7 | Pf04352 |
| 4.49E+04 | FTL_0476 | | 81.9 | Pf01276, Pf03709, Pf03711 |
| 4.03E+04 | FTL_0075 | ribH | 16.3 | Pf00885 |
| 3.99E+04 | FTL_1659 | prfC | 59.4 | Pf00009, Pf03144, Pf16658 |
| 1.94E+04 | FTL_0822 | | 9.6 | |
| 1.26E+03 | FTL_0241 | rplV | 12.2 | Pf00237 |

**Table S2.** Strains and plasmids used in this study.

| Strain or Plasmid | Characteristics[a] | Source |
|---|---|---|
| *F. tularensis* Live Vaccine Strain (LVS) | *Francisella tularensis* subsp. *holartica* Live Vaccine Strain (LVS) | ATCC |
| LVS Δ*pilE4* | LVS with deletion of *pilE4* | This study |
| LVS Δ*pilF* | LVS with deletion of *pilF* | This study |
| LVS Δ*pilT* | LVS with deletion of *pilT* | This study |
| LVS *att*::*NLL-MetRS* | LVS with the *NLL-MetRS* modified methionyl-tRNA synthetase (1) inserted into the chromosomal Tn7 *att* site, $Kan^R$ | This study |
| LVS *ΔpilE4 att*::*NLL-MetRS* | LVS *ΔpilE4* with *NLL-MetRS* inserted into the chromosomal Tn7 *att* site, $Kan^R$ | This study |
| LVS *ΔpilF att*::*NLL-MetRS* | LVS *ΔpilF* with *NLL-MetRS* inserted into the chromosomal Tn7 *att* site, $Kan^R$ | This study |
| LVS *ΔpilT att*::*NLL-MetRS* | LVS *ΔpilT* with *NLL-MetRS* inserted into the chromosomal Tn7 *att* site, $Kan^R$ | This study |
| LVS *att*::*chiA-3xFLAG* | LVS with insertion of *chiA-3xFLAG* into the chromosomal Tn7 *att* site, $Kan^R$ | This study |
| LVS *ΔpilE4 att*::*chiA-3xFLAG* | LVS *ΔpilE4* with insertion of *chiA-3xFLAG* into the chromosomal Tn7 *att* site, $Kan^R$ | This study |
| LVS *ΔpilF att*::*chiA-3xFLAG* | LVS *ΔpilF* with insertion of *chiA-3xFLAG* into the chromosomal Tn7 *att* site, $Kan^R$ | This study |
| LVS *ΔpilT att*::*chiA-3xFLAG* | LVS *ΔpilT* with insertion of *chiA-3xFLAG* into the chromosomal Tn7 *att* site, $Kan^R$ | This study |
| LVS *att*::*chiD-3xFLAG* | LVS with insertion of *chiD-3xFLAG* into the chromosomal Tn7 *att* site, $Kan^R$ | This study |
| LVS *ΔpilE4 att*::*chiD-3xFLAG* | LVS *ΔpilE4* with insertion of *chiD-3xFLAG* into the chromosomal Tn7 *att* site, $Kan^R$ | This study |
| LVS *ΔpilF att*::*chiD-3xFLAG* | LVS *ΔpilF* with insertion of *chiD-3xFLAG* into the chromosomal Tn7 *att* site, $Kan^R$ | This study |
| LVS *ΔpilT att*::*chiD-3xFLAG* | LVS *ΔpilT* with insertion of *chiD-3xFLAG* into the chromosomal Tn7 *att* site, $Kan^R$ | This study |

| | | |
|---|---|---|
| LVS *att*::*katG-3xFLAG* | LVS with insertion of *katG-3xFLAG* into the chromosomal Tn7 *att* site, $Kan^R$ | This study |
| LVS *ΔpilE4 att*::katG-*3xFLAG* | LVS *ΔpilE4* with insertion of *katG-3xFLAG* into the chromosomal Tn7 *att* site, $Kan^R$ | This study |
| LVS *ΔpilF att*::*katG-3xFLAG* | LVS *ΔpilF* with insertion of *katG-3xFLAG* into the chromosomal Tn7 *att* site, $Kan^R$ | This study |
| LVS *ΔpilT att*::*katG-3xFLAG* | LVS *ΔpilT* with insertion of *katG-3xFLAG* into the chromosomal Tn7 *att* site, $Kan^R$ | This study |
| LVS *att*::*sucC-3xFLAG* | LVS with insertion of *sucC-3xFLAG* into the chromosomal Tn7 *att* site, $Kan^R$ | This study |
| LVS *ΔpilE4 att*::*sucC-3xFLAG* | LVS *ΔpilE4* with insertion of *sucC-3xFLAG* into the chromosomal Tn7 *att* site, $Kan^R$ | This study |
| LVS *ΔpilF att*::*sucC-3xFLAG* | LVS *ΔpilF* with insertion of *sucC-3xFLAG* into the chromosomal Tn7 *att* site, $Kan^R$ | This study |
| LVS *ΔpilT att*::*sucC-3xFLAG* | LVS *ΔpilT* with insertion of *sucC-3xFLAG* into the chromosomal Tn7 *att* site, $Kan^R$ | This study |
| LVS *att*::*FTL_0569-3xFLAG* | LVS with insertion of *FTL_0569-3xFLAG* into the chromosomal Tn7 *att* site, $Kan^R$ | This study |
| LVS *ΔpilE4 att*::*FTL_0569-3xFLAG* | LVS *ΔpilE4* with insertion of *FTL_0569-3xFLAG* into the chromosomal Tn7 *att* site, $Kan^R$ | This study |
| LVS *ΔpilF att*::*FTL_0569-3xFLAG* | LVS *ΔpilF* with insertion of *FTL_0569-3xFLAG* into the chromosomal Tn7 *att* site, $Kan^R$ | This study |
| LVS *ΔpilT att*::*FTL_0569-3xFLAG* | LVS *ΔpilT* with insertion of *FTL_0569-3xFLAG* into the chromosomal Tn7 *att* site, $Kan^R$ | This study |
| LVS *att*::*accB-3xFLAG* | LVS with insertion of *accB-3xFLAG* into the chromosomal Tn7 *att* site, $Kan^R$ | This study |
| LVS *ΔpilE4 att*::*accB -3xFLAG* | LVS *ΔpilE4* with insertion of *accB-3xFLAG* into the chromosomal Tn7 *att* site, $Kan^R$ | This study |
| LVS *ΔpilF att*::*accB -3xFLAG* | LVS *ΔpilF* with insertion of *accB-3xFLAG* into the chromosomal Tn7 *att* site, $Kan^R$ | This study |
| LVS *ΔpilT att*::*accB-3xFLAG* | LVS *ΔpilT* with insertion of *accB-3xFLAG* into the chromosomal Tn7 *att* site, $Kan^R$ | This study |
| LVS *att*::*FTL_1225-3xFLAG* | LVS with insertion of *FTL_1225-3xFLAG* into the chromosomal Tn7 *att* site, $Kan^R$ | This study |

| | | |
|---|---|---|
| LVS *ΔpilE4 att*::*FTL_1225-3xFLAG* | LVS *ΔpilE4* with insertion of *FTL_1225-3xFLAG* into the chromosomal Tn7 *att* site, $Kan^R$ | This study |
| LVS *ΔpilF att*::*FTL_1225-3xFLAG* | LVS *ΔpilF* with insertion of *FTL_1225-3xFLAG* into the chromosomal Tn7 *att* site, $Kan^R$ | This study |
| LVS *ΔpilT att*::*FTL_1225-3xFLAG* | LVS *ΔpilT* with insertion of *FTL_1225-3xFLAG* into the chromosomal Tn7 *att* site, $Kan^R$ | This study |
| pMP812 | Plasmid for generation of chromosomal deletion mutants by allelic exchange, $Kan^R$ | (2) |
| pMP749 | Mini-Tn7 suicide vector for stable integration of insert into the chromosomal *att* site, $Kan^R$ | (2) |
| pMP720 | Helper plasmid for generation of chromosomal insertions using plasmid pMP749, $Hyg^R$ | (2) |
| pMP831 | Shuttle vector for plasmid-based gene expression in *F. tularensis*, $Hyg^R$ | (3) |
| pMP822 | Shuttle vector for plasmid-based gene expression in *F. tularensis*, with *blaB* overexpression promoter, $Hyg^R$ | (3) |
| pMP749-NLL-MetRS | pMP749 containing the *NLL-MetRS* modified methionyl-tRNA synthetase (1), $Kan^R$ | This Study |
| pMP749-chiA-3xFLAG | pMP749 containing *chiA-3x* FLAG, $Kan^R$ | This Study |
| pMP749-chiD-3xFLAG | pMP749 containing *chiD-3x* FLAG, $Kan^R$ | This Study |
| pMP749-katG-3xFLAG | pMP749 containing k*atG-3x* FLAG, $Kan^R$ | This Study |
| pMP749-sucC-3xFLAG | pMP749 containing *sucC-3x* FLAG, $Kan^R$ | This Study |
| pMP749-AccB-3xFLAG | pMP749 containing AccB-3*x* FLAG, $Kan^R$ | This Study |
| pMP749-FTL_0569-3xFLAG | pMP749 containing FTL_0569 *3x* FLAG, $Kan^R$ | This Study |
| pMP749-FTL_1225-3xFLAG | pMP749 containing FTL_1225 3x FLAG, $Kan^R$ | This Study |
| pMP831-pilE4 | pMP831 containing LVS *pilE4* under its natural promoter, $Hyg^R$ | This study |
| pMP831-pilF | pMP831 containing LVS *pilF* under its natural promoter, $Hyg^R$ | This study |

| | | |
|---|---|---|
| pMP831-pilT$_{LVS}$ | pMP831 containing LVS *pilT* under its natural promoter, Hyg$^R$ | This study |
| pMP831-pilT$_{F.n.}$ | pMP831 containing *F. novicida pilT* under its natural promoter, Hyg$^R$ | This study |
| pMP822-pilT$_{LVS}$ | pMP822 containing LVS *pilT* under the *blaB* over-expression promoter, Hyg$^R$ | This study |
| pMP822-pilT$_{F.n.}$ | pMP822 containing *F. novicida pilT* under the *blaB* over-expression promoter, Hyg$^R$ | This study |

[a]Kan$^R$, kanamycin resistance; Hyg$^R$, hygromycin resistance.

**Table S3.** Primers used in this study.

| Primer Name | Nucleotide Sequence (5`→3`) | Description |
|---|---|---|
| LVSdelPilE4_up_F | GCGCGCGGCCGCTAATAACAACCGTAATAATAATGCTGCT | Upstream region for *pilE4* deletion (forward) |
| LVSdelPilE4_up_R | GGGGGCCCCCGGGGGCAAAAATGCTTGGTCTTGCTAAAAA | Upstream region for *pilE4* deletion (reverse) |
| LVS_delPilE4_down_F | CCCCCGGGGGCCCCCATCACTCCCGGAAAAATTGTTTAAA | Downstream region for *pilE4* deletion (forward) |
| LVS_delPilE4_down_R | GCGCGTCGACCAAAAATGCTTGGTCTTGCTAAAAA | Downstream region for *pilE4* deletion (reverse) |
| LVS_delPilF_up_F | GCGCGGATCCAAACATCATTCTCACTAGCC | Upstream region for *pilF* deletion (forward) |
| LVS_delPilF_up_R | TTCCCTTTAACTTACACGGTCTATATAAAATAAACTTCAAATAGTAGCA | Upstream region for *pilF* deletion (reverse) |
| LVS_delPilF_down_F | TGCTACTATTTGAAGTTTATTTATATAGACCGTGTAAGTTAAAGGGAA | Downstream region for *pilF* deletion (forward) |
| LVS_delPilF_down_R | GCGCGTCGACTCCCTGATTCTTCACCAGCAGT | Downstream region for *pilF* deletion (reverse) |
| PilT upstream_F | GCGCGGATCCTCTTCACGTGACTGGGCTGC | Upstream region for *pilT* deletion (forward) |
| PilT_upstream_R | GCTAGCAATATAAATATTAGATTTTTAAATGAATCAAGAATAGAACTTAAATTTAATAATAG | Upstream region for *pilT* deletion (reverse) |
| PilT_downstream_F | TATTATTAAATTTAAGTTCTATTCTTGATTCATTTAAAAATCTAATATTTATATTGCTAGC | Downstream region for *pilT* deletion (forward) |
| PilT_downstream_R | GCGCGTCGACACCAATGGCTTGTCAATGATGA | Downstream region for *pilT* deletion (forward) |
| LVS_ChiA_3xFLAG_F1 | GAGACGGTACCATAGGTTCTTGGGGTGTTGC | Forward primer for amplification of *chiA* |
| LVS_ChiA_3xFLAG_R1 | GTGGTCCTTGTAGTCTTGTTTTTCCCAAACATT | Reverse primer for amplification of *chiA* with 3xFLAG overlap |
| LVS_ChiA_3xFLAG_F2 | AATGTTTGGGAAAAACAAGACTACAAGGACCAC | Forward primer for amplification of *chiA* with 3xFLAG overlap |

| | | |
|---|---|---|
| LVS_ChiD_3xFLAG_F1 | GAGACGGTACCGCTATTATCTACTTTACCGGC | Forward primer for amplification of *chiD* |
| LVS_ChiD_3xFLAG_R1 | GTGGTCCTTGTAGTCTTTACTATCTATTTTTGTCCA | Reverse primer for amplification of *chiD* with 3xFLAG overlap |
| LVS_ChiD_3xFLAG_F2 | TGGACAAAAATAGATAGTAAAGACTACAAGGACCAC | Forward primer for amplification of *chiD* with 3xFLAG overlap |
| FTL_1592-3xFLAG_F1 | GCGCGGTACCATGTGTGTTTGGAGGTAAGC | Forward primer for amplification of FTL_1592 |
| FTL_1592_3xFLAG_R1 | GTGGTCCTTGTAGTCTTCAATAATAAATAGAGG | Reverse primer for amplification of FTL_1592 with 3xFLAG overlap |
| FTL_1592-3xFLAG_F2 | CCTCTATTTATTATTGAAGACTACAAGGACCAC | Forward primer for amplification of FTL_1592 with 3xFLAG overlap |
| LVS_SucC_3x_F1 | GAGACGGTACCTGTACCAAACTGGATTGC | Forward primer for amplification of *sucC* |
| LVS_SucC_3xFLAG_R1 | GTGGTCCTTGTAGTCACCTAGTGATTTCACAAC | Reverse primer for amplification of *sucC* with 3xFLAG overlap |
| LVS_SucC_3xFLAG_F2 | GTTGTGAAATCACTAGGTGACTACAAGGACCAC | Forward primer for amplification of *sucC* with 3xFLAG overlap |
| FTL_1225_3xFLAG_F1 | GAGACGGTACCCTAAAATTAGTTGACGTTGATGC | Forward primer for amplification of FTL_1225 |
| FTL_1225_3xFLAG_R1 | GTGGTCCTTGTAGTCTTTCTCCATAAATGTAAC | Reverse primer for amplification of FTL_1225 with 3xFLAG overlap |
| FTL_1225_3xFLAG_F2 | GTTACATTTATGGAGAAAGACTACAAGGACCAC | Forward primer for amplification of FTL_1225 with 3xFLAG overlap |

| | | |
|---|---|---|
| LVS_KatG_3xFLAG_F1 | GCGCGGGTCCTAGAACCTACCCCTATACCAAC | Forward primer for amplification of *katG* |
| LVS_KatG_3xFLAG_R1 | GTGGTCCTTGTAGTCGCCAAGCATCATAACTT | Reverse primer for amplification of *katG* with 3xFLAG overlap |
| LVS_KatG_3xFLAG_F2 | AAGTTATGATGCTTGGCGAC TAC AAG GAC CAC | Forward primer for amplification of *katG* with 3xFLAG overlap |
| NLL-MetRS_rpsl | GGAGAATAATAACTAATGACTCAAGTCGCGAAGAAAATTCTGGTG | Forward primer for amplification of open reading frame of NLL-metRS |
| NLL-MetRS_bamHI | GCGCGCGGATCCTCATTTAGAGGCTTCCACCAGTGC | Reverse primer for amplification of open reading frame of NLL-metRS |
